# CTCF Mediates Dosage and Sequence-context-dependent Transcriptional Insulation through Formation of Local Chromatin Domains

**DOI:** 10.1101/2020.07.07.192526

**Authors:** Hui Huang, Quan Zhu, Adam Jussila, Yuanyuan Han, Bogdan Bintu, Colin Kern, Mattia Conte, Yanxiao Zhang, Simona Bianco, Andrea Chiariello, Miao Yu, Rong Hu, Ivan Juric, Ming Hu, Mario Nicodemi, Xiaowei Zhuang, Bing Ren

## Abstract

Insulators play a critical role in spatiotemporal gene expression in metazoans by separating active and repressive chromatin domains and preventing inappropriate enhancer-promoter contacts. The evolutionarily conserved CCCTC-binding factor (CTCF) is required for insulator function in mammals, but not all of its binding sites act as insulators. Here, we explore the sequence requirements of CTCF-mediated transcriptional insulation with the use of a sensitive insulator reporter assay in mouse embryonic stem cells. We find that insulation potency depends on the number of CTCF binding sites in tandem. Furthermore, CTCF-mediated insulation is dependent on DNA sequences flanking its core binding motifs, and CTCF binding sites at topologically associating domain(TAD) boundaries are more likely to function as insulators than those outside TAD boundaries, independent of binding strength. Using chromosomal conformation capture assays and high-resolution chromatin imaging techniques, we demonstrate that insulators form local chromatin domain boundaries and reduce enhancer-promoter contacts. Taken together, our results provide strong genetic, molecular, and structural evidence connecting chromatin topology to the action of insulators in the mammalian genome.

## Introduction

The spatial and temporal patterns of gene expression are encoded in the genome sequences in the form of cis-regulatory elements which are categorized into promoters, enhancers, insulators, and other less-studied regulatory sequences, including repressive/silencing elements^1-3^. In metazoans, insulators play an essential role in cell-type-specific gene expression by protecting genes from improper regulatory signals from the neighboring chromatin environment^4^. One class of insulators acts as barriers to heterochromatin spreading^5^, while the other blocks enhancer-promoter communications^6^. Enhancer-blocking (EB) insulators act in a position-dependent manner in that they prevent enhancer-dependent gene activation only when placed in between the enhancer and target gene^6-8^. Insulators were initially identified in *Drosophila*, where the molecular machinery for insulation was first elucidated^4, 6, 9^. The first identified enhancer-blocking insulator in vertebrates is the 5’-HS4 element of the chicken β-globin locus^10^. Detailed analysis of this insulator led to the finding that the evolutionarily conserved zinc-finger family transcription factor CTCF, first identified as DNA binding protein at the chicken *c-Myc* gene promoter^11^, was essential for its enhancer-blocking activity^12^. Mutations in the CTCF protein or its binding sites at insulators have since been implicated in a broad spectrum of human diseases^13-15^. In addition to its function at insulators, CTCF has also been demonstrated to play roles in transcriptional repression, gene activation, alternative splicing, and class switch recombination depending on the context of genomic locus^11, 16-20^. There are reports that CTCF binding at gene promoters could promote, instead of block, enhancer-promoter interactions^21, 22^. To date, exactly how and where CTCF mediates insulator function remains unclear.

CTCF has long been postulated to function as an organizer of the three-dimensional chromosome architecture^1, 23, 24^. Genome-wide chromosome conformation capture analyses showed that the interphase chromosomes in mammalian cells are partitioned into megabase-sized topologically associating domains (TADs)^25, 26^, and the binding sites for CTCF were found at over 75% of TAD boundaries^25^, suggesting a probable link between TAD boundaries and CTCF-mediated transcriptional insulation. Supporting this connection, disruption of TAD boundaries has been shown to permit ectopic enhancer-promoter contacts and aberrant gene expression, thereby leading to developmental abnormalities and cancer^17, 27^. Additionally, depletion of CTCF can lead to the weakening or disappearance of TADs^28-30^. CTCF drives TAD formation by working together with the cohesin complex to establish dynamic chromatin loops between distant CTCF binding sites, likely through a loop-extrusion process^30-40^ or other mechanisms such as phase separation^41-46^. However, it is still debated whether TAD boundaries are sufficient to provide transcriptional insulation. Rapidly dissolving the global TAD structure by acute depletion of CTCF or cohesin subunits only altered transcription of a small number of genes in many different cellular contexts^28, 30, 34, 36, 38, 47^. Moreover, deletion of CTCF sites at the developmental gene *Sox9-Kcnj2* TAD boundary did not cause discernible phenotypes^48^. Furthermore, a majority of CTCF binding sites are not located at TAD boundaries, and whether these CTCF sites may function as insulators is unclear. These observations warrant an in-depth investigation of the role that CTCF and TADs play in transcriptional insulation.

To better understand where and how CTCF may mediate transcriptional insulation in the genome, we have developed an insulator reporter assay to evaluate the function of any DNA fragments in blocking enhancer-dependent transcriptional activation in mouse embryonic stem (mES) cells. Using this system, we demonstrated that isolated single CTCF sites have weak or no insulator activity, regardless of its DNA binding strength as measured via biochemical assays. Instead, multiple copies of CTCF sites placed in tandem can provide a potent insulation effect. We also observed that CTCF binding sites at TAD boundaries could function as potent insulators, while the CTCF sites not located at TAD boundaries were incapable of insulating transcription. We attributed this difference in insulation activity to sequences immediately flanking the CTCF core motifs. We further discovered that insulators act by forming local TAD boundaries to reduce long-range enhancer-promoter contacts, using both chromosome conformation capture assays and high throughput multiplexed DNA fluorescence *in situ* hybridization (FISH) techniques. These results, taken together, shed new light on how CTCF mediates transcriptional insulation in mammalian cells and establish a direct link between TAD boundaries and insulators.

## Results

### A sensitive insulator reporter assay in mouse embryonic stem cells

To quantitatively assay insulator activities in the context of native chromatin in cells, we engineered the *Sox2* gene locus in the F123 mES cell line, which was derived from a hybrid F1 mouse progeny (*Mus musculus castaneus* × *S129/SvJae*)^49^. We and others previously showed that a super-enhancer located ∼110kb downstream of the *Sox2* gene was responsible for over 90% of its expression in the mES cells ^50, 51^. We reasoned that insulator activity of DNA elements could be measured by the reduction in *Sox2* gene expression when inserted between the *Sox2* gene and the downstream super-enhancer. Therefore, we first tagged the two copies of the *Sox2* gene with *egfp (*CAST allele) and *mCherry (*129 allele) to quantify allelic Sox2 expression by live-cell fluorescence-activated cell sorting (FACS) (Fig. 1a, Extended Data Fig. 1a). Subsequently, we inserted a suicidal fusion gene Tg(CAG-*HyTK*) flanked by a pair of heterotypic Flippase recognition sites (*Frt/F3*) between the *Sox2* gene and its downstream super-enhancer (*SE*) on the CAST allele (Fig. 1a, Extended Data Fig. 1b). As enhancer-blocking insulation is position-dependent, we created a control clone with the same replaceable cassette placed further downstream of the *Sox2* super-enhancer at equal distance on the CAST allele (Fig. 1a, Extended Data Fig. 1c). The suicidal marker gene can be replaced by a donor sequence using the **<u>r</u>**ecombinase **<u>m</u>**ediated **<u>c</u>**assette **<u>e</u>**xchange (RMCE) strategy (Fig. 1b, Extended Data Fig. 2a). By killing off unmodified mES cells with ganciclovir, we could achieve nearly 100% efficiency of marker-free insertion (Extended Data Fig. 2b).

**Fig. 1.**
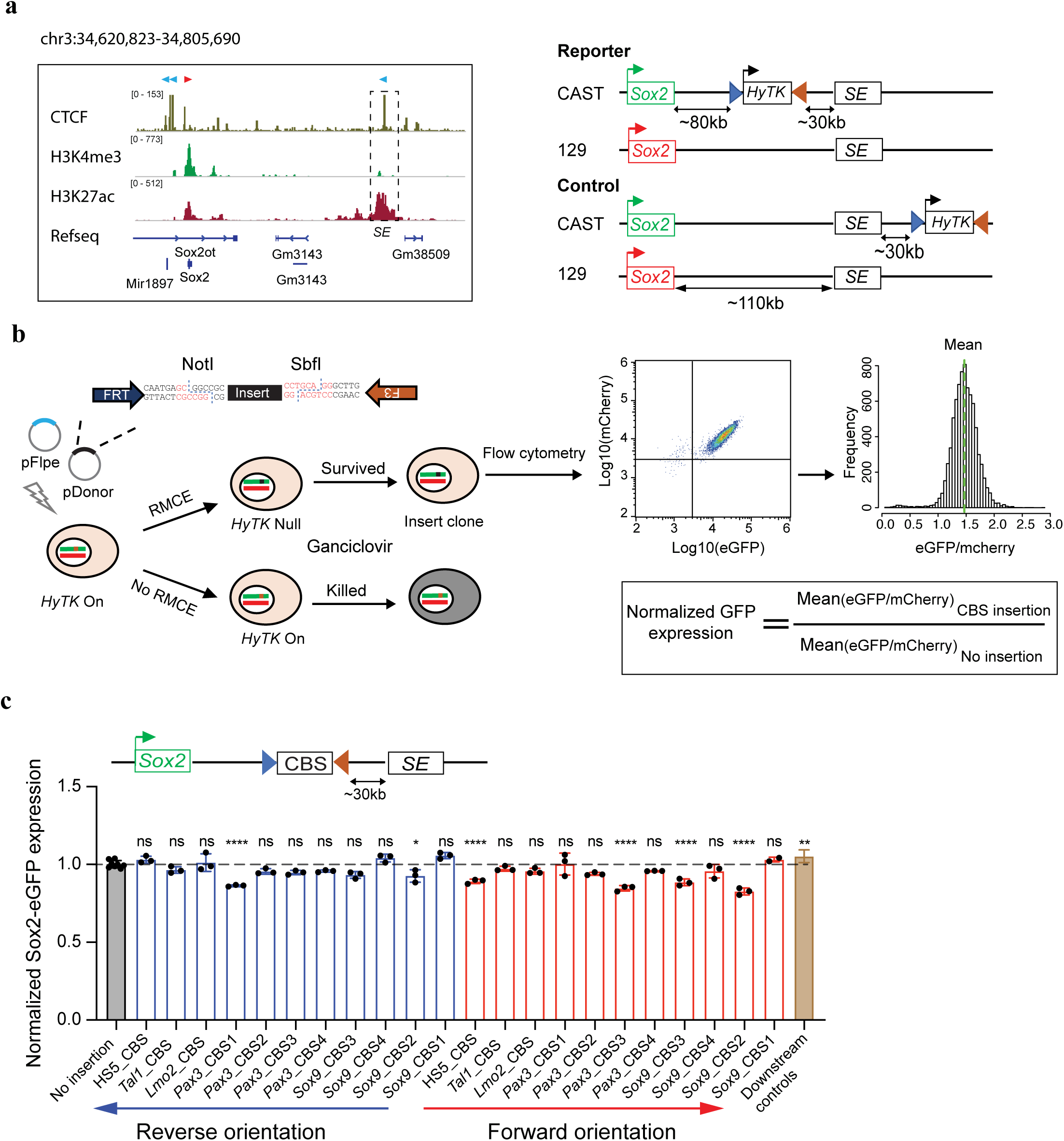
A sensitive insulator reporter assay measures the insulation activity of different CTCF binding sites at the *Sox2* locus in mouse ES cells. **a**, Left, the regulatory landscape of the *Sox2* locus in mES cells. Orientations of CTCF sites are indicated on the top of the signal tracks; Right, genetic constructs of mES cell lines. Boxed *Sox2* in green represents *Sox2-p2a-egfp in situ* fusion gene, boxed *Sox2* in red represents *Sox2-p2a-mCherry in situ* fusion gene. The hygromycin phosphotransferase-thymidine kinase fusion gene *HyTK* is flanked by Flippase recognition sites *FRT* and *F3*. **b**, Experimental scheme to insert a test sequence into the *Sox2* locus by recombinase mediated cassette exchange (RMCE). The Flippase expression plasmid and donor plasmid containing the test sequence were co-electroporated into cells. The donor plasmid contains Not1 and Sbf1 restriction enzyme sites so that the orientation of the insert can be controlled. Mouse ES cell clones containing the insert were picked, genotyped, and allelic Sox2 expression was measured by FACS. **c**, A bar graph shows the normalized Sox2-eGFP expression of the no insertion clone (n=8), different CBS insertion clones (n=3. For Sox9_CBS1 in forward orientation, n=2.) and downstream insertion controls (n=27). Each dot represents an independently picked colony. One-way analysis of variance with Bonferroni’s multiple comparisons test. ns *P* > 0.05, \**P* ≤ 0.05, \*\**P* ≤ 0.01, \*\*\**P* ≤ 0.001, \*\*\*\**P* ≤ 0.0001. Data are mean ± sd.

As the insertion was specifically on the CAST allele, we used the 129 allele as the internal control to correct clone-to-clone variations in Sox2 expression (Fig. 1b, Extended Data Fig. 3a-b), which allowed quantitative comparisons of insulator activities of different CTCF binding sites (CBSs). We tested the insulation activity of a total of 11 different CBSs selected from several known TAD boundaries and chromatin loop anchors (Supplementary Table1). Each CBS insert was amplified from mouse or human genomic DNA by PCR and was 1-4kb in length. Surprisingly, isolated single CBS tested in both the forward and reverse orientations generally exhibited little or no insulator effect (Fig. 1c). Only two of the probed CBSs in reverse orientation and four of the probed CBSs in forward orientation showed significant yet modest insulator effects (Fig. 1c). The CBS of a canonical insulator, the HS5 sequence of the human beta-globin locus, reduced Sox2 expression by 11.0%± 1.9% when inserted in forward orientation but had no effect in reverse orientation (Fig. 1c, Extended Data Fig. 3c-d). On average, individual isolated CBS in forward and reverse orientations reduced Sox2 expression to 93.0%(+/-6.5%) and 97.0%(+/-6.0%) of parental cells with no insertion, respectively (Fig. 1c).

### Multiple CTCF sites in tandem enable strong transcriptional insulation

Since single CBS was weak in transcriptional insulation, we hypothesized that multiple CBSs collectively may provide more robust insulation, given that TAD boundaries are enriched for clustered CTCF binding sites^25, 52^. To test this possibility, we constructed a series of insertion clones harboring multiple CBSs from the *Sox9-Kcnj2* TAD boundary (Extended Data Fig. 4a). Two or more CBSs were PCR-amplified from mouse genomic DNA, ligated together and inserted in between the *Sox2* gene and *SE* on the CAST allele by RMCE as described above. We found that two CBSs, in forward tandem, reverse tandem, or divergent orientations, all had significantly stronger insulation effect than individual CBSs alone (Fig. 2a). Notably, combining a weak CBS insulator with one that had a negligible insulator activity gave rise to stronger insulation than the summed effects of the two individual sites (Fig. 2a), suggesting that CBSs could have synergistic insulation effects. Next, we measured the insulator activity of CBS clusters consisting of up to all four CBSs from the *Sox9-Kcnj2* TAD boundary. ChIP-seq analysis indicated that CTCF was recruited to the extra copy of the boundary sequence inserted in the *Sox2* domain (Extended data Fig. 4b). We found that the insulation effect became stronger as the number of CBS increased, regardless of the orientation of CTCF motifs (Fig. 2b). Interestingly, the enhancement of insulation conferred by each additional CBS became smaller when the number of CBSs exceeds two (Extended Data Fig. 4c).

**Fig. 2.**
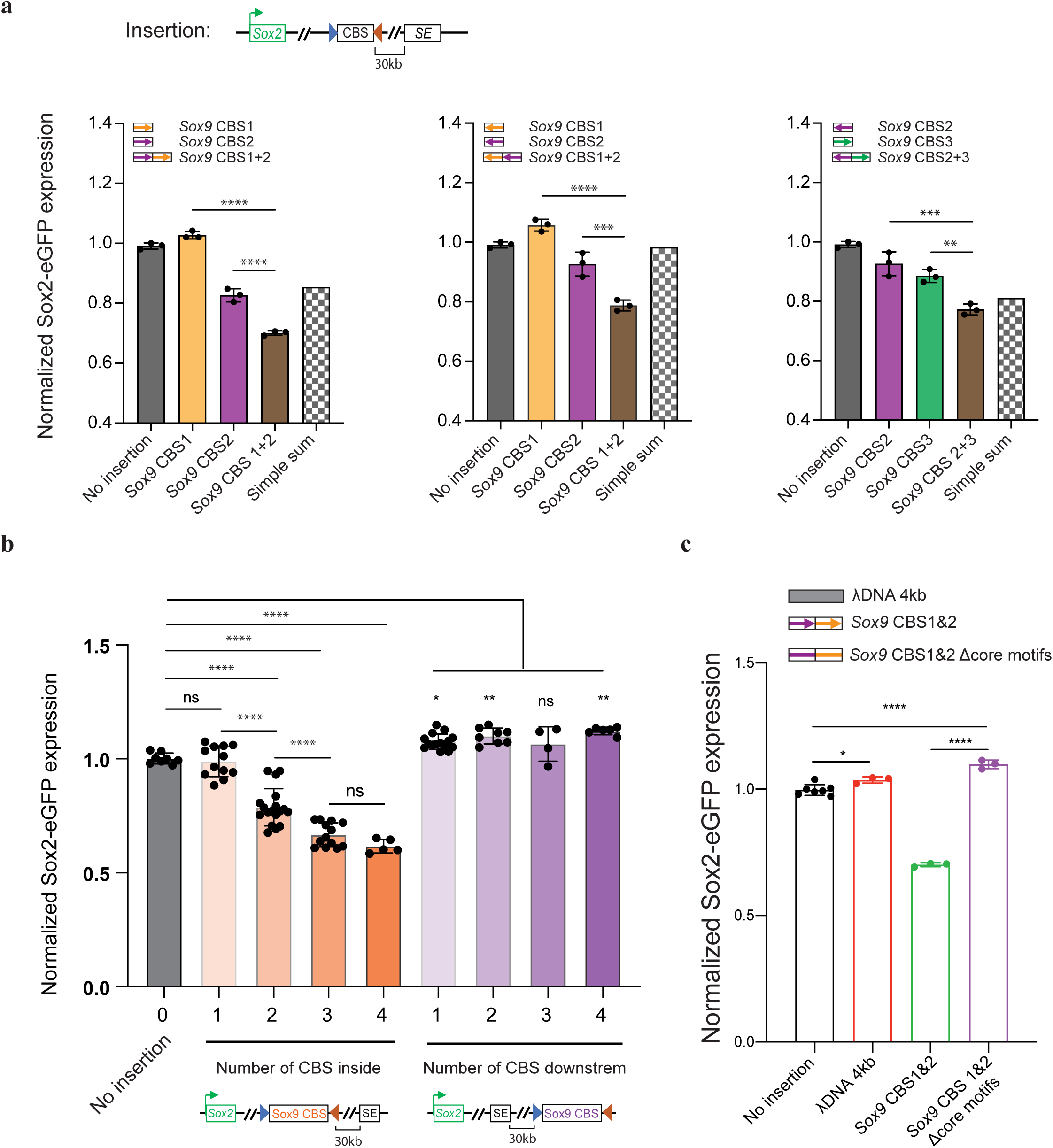
Multiple CTCF sites in tandem enable strong transcriptional insulation. **a**, A bar graph shows additive or synergistic insulation effects by two CBSs from the *Sox9-Kcnj2* TAD boundary (n=3). Individual CBS sequences were combined by PCR to create two-CBS insertions. Arrows indicate motif orientation of every CBS. Every insertion construct was created by an independent RMCE experiment. **b**, A bar graph shows insulation effects of multiple CBS from the *Sox9-Kcnj2* TAD boundary. Individual or combined CBS sequences were PCR cloned from mouse genomic DNA. Motif orientations of CBSs were kept the same as in the *Sox9-Kcnj2* TAD boundary. Each dot represents an independent clone created by RMCE. 0 CBS, n=8; 1 CBS inside, n=12; 2 CBS inside, n=18; 3 CBS inside, n=13; 4 CBS inside, n=5; 1 CBS downstream, n=15; 2 CBS downstream, n=8; 3 CBS downstream, n=4; 4 CBS downstream, n=6. **c**, A bar graph shows insulation effects of λ DNA (n=3), a combined two-CBS sequence, *Sox9* CBS1&2 (n=3), and *Sox9* CBS1&2 Δcore motifs, which is the same 2-CBS sequence but with the two19-bp CTCF core motifs deleted (n=3). Inserts were comparable in length (∼4kb). One-way analysis of variance with Bonferroni’s multiple comparisons test. ns *P* > 0.05, \**P* ≤ 0.05, \*\**P* ≤ 0.01, \*\*\**P* ≤ 0.001, \*\*\*\**P* ≤ 0.0001. Data are mean ± sd.

Consistent with the requirement for CTCF in transcriptional insulation, removal of the binding motifs of CTCF within the inserts completely abolished insulation effects of CBSs (Fig. 2c). Furthermore, introducing CTCF sites downstream of the *Sox2 SE* did not reduce but rather slightly increased Sox2 expression, likely due to insulation of interactions between the *SE* and further downstream chromatin (Fig. 2b). Taken together, these results suggest that multiple CTCF binding sites arranged in tandem can function as a potent insulator due to synergistic or additive effects from individual sites.

Surprisingly, we observed that the insulator containing four CBSs was able to reduce Sox2 expression by 38.47 ± 3.16%, rather than completely blocking the *Sox2 SE* activity. Interestingly, this insulator substantially increased cell-to-cell variations in Sox2 expression, evidenced by the accumulation of cells with extremely low Sox2-eGFP signals (Extended Data Fig. 4d). Moreover, the sub-population of cells expressing ultra-low Sox2-eGFP could revert to the state of higher expression level after extended culturing, suggesting that the cell-to-cell variation of *Sox2* gene expression was a meta-stable state (Extended Data Fig. 4e). Furthermore, CTCF insulation did not change the active chromatin state on either the *Sox2* promoter or its enhancer (Extended data Fig. 4f-g). Collectively, these results suggest that CBS-mediated insulation is permissive and highly dynamic.

### CTCF-mediated insulator function depends on sequence contexts

To better understand the sequence requirements for CTCF-mediated insulation, we synthesized insulators by concatenating multiple 139-bp genomic DNA sequences, each containing a 19-bp CTCF motif at the center surrounded by two 60-bp flanking sequences. Each site was selected from the aforementioned CBSs (four CBSs from the *Sox9-Kcnj2* TAD boundary, one from the *Pax3-Epha4* TAD boundary and one from the human β-globin HS5 CBS, Supplementary Table 2). Consistent with observations described above, the synthetic DNA sequences showed additive effects in transcriptional insulation (Extended Data Fig. 5a). Additionally, ChIP-seq analysis confirmed the recruitment of CTCF and the cohesin complex to the synthetic insulators (Fig. 3a). Interestingly, we observed that CBSs with longer flanking sequences (1-kb or longer) had stronger insulation effects than the shorter 139-bp CBSs, suggesting the existence of additional elements that could facilitate insulation (Extended Data Fig. 5b).

**Fig. 3.**
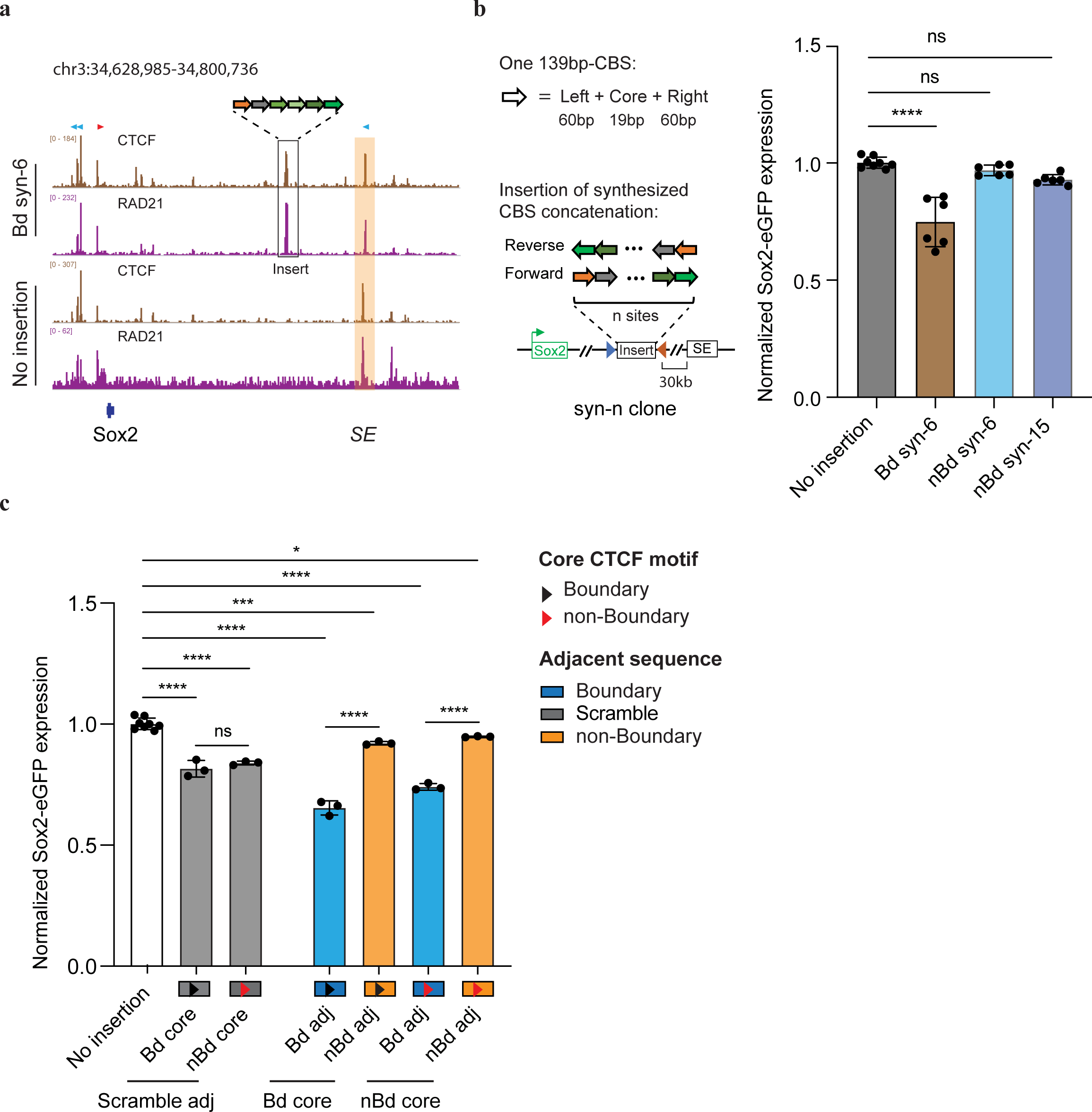
Synthetic insulators reveal sequence requirements for CTCF-mediated enhancer-blocking. **a**, ChIP-seq of CTCF and Rad21. The “Bd syn-6” mES clone contains the insertion of six 139-bp boundary CBS (four *Sox9-Kcnj2* boundary CBSs, one *Pax3-Epha4* boundary CBS and the human β-globin HS5 CBS) between *Sox2* and its super-enhancer. Sequencing reads from no insertion cells were aligned to the mm10 reference genome. Sequencing reads from the insertion clone were aligned to a customized mm10 genome that included the inserted sequence at the target location. Motif orientations of nearby CBS and inserted CBS were indicated on the top of signal tracks. The *Sox2* super-enhancer is highlighted in the orange box. **b**, A bar plot shows insulation effects of synthetic sequences containing tandemly arrayed 139bp-CBS from boundary and non-boundary regions. Synthetic sequences were inserted between *Sox2* and its super-enhancer. For each synthetic sequence, six insertion clones were picked with three of them in forward orientation and the other three in reverse orientation (n=6). One-way analysis of variance with Bonferroni’s multiple comparisons test. ns *P* > 0.05, \**P* ≤ 0.05, \*\**P* ≤ 0.01, \*\*\**P* ≤ 0.001, \*\*\*\**P* ≤ 0.0001. Data are mean ± sd. **c**, A bar plot shows insulation effects of recombined tandemly arrayed 139bp-CBS. CBS core motifs of boundary and non-boundary sites were combined with either their native adjacent sequences, scrambled adjacent sequences, or exchanged adjacent sequences with each other (n=3). Each test sequence contains six tandemly arrayed 139bp-CBS. The order of the six CBS core motifs was kept the same. One-way analysis of variance with Bonferroni’s multiple comparisons test. ns *P* > 0.05, \**P* ≤ 0.05, \*\**P* ≤ 0.01, \*\*\**P* ≤ 0.001, \*\*\*\**P* ≤ 0.0001. Data are mean ± sd.

Using the same approach, we also tested whether CBSs from outside of TAD boundaries could function as insulators. We selected multiple CBSs from non-TAD boundary regions in the genome, concatenated multiple 139-bp genomic sequences containing CTCF binding motifs together, and tested their insulation ability in our insulator reporter assay (Supplementary Table 3). Surprisingly, although these non-TAD boundary CBSs displayed stronger CTCF binding than those from TAD boundaries at their original loci, the synthetic DNA sequences made up of six or fifteen tandemly arrayed 139-bp CBSs from non-boundary regions were unable to function as insulators, despite presence of strong CTCF ChIP-seq signals (Fig 3b, Extended Data Fig. 5c-d), indicating that CTCF binding alone is insufficient to bring transcriptional insulation.

To further dissect the sequence dependence of CTCF-mediated insulation, we exchanged the core motifs of 139-bp boundary CBSs with those of the synthetic CBSs from non-boundary regions. Combining boundary CBS core motifs with non-boundary adjacent sequences resulted in a much weaker insulation effect than with their original neighboring sequences of equal lengths (Fig. 3c). In contrast, replacing adjacent sequences of non-boundary CBSs with those from boundary sites significantly strengthened their insulation effect (Fig. 3c). However, when the adjacent sequences were scrambled or kept the same for boundary and non-boundary core motifs, their effects in insulating Sox2 expression were comparable (Fig. 3c). Together, these results suggest that transcriptional insulation by CTCF is sequence-context-dependent, requiring DNA elements flanking the CTCF binding motif.

### Insulators promote formation of local chromatin domains and reduce enhancer-promoter contacts

Previous data suggest that the *Sox2 SE* forms long-range chromatin contacts with the *Sox2* promoter^51, 53^. We hypothesized that insulators may change chromosome topology to limit enhancer-promoter communication. To test this hypothesis, we performed PLAC-seq^54^ (also known as HiChIP^55^) experiments using mES cell clones with various insulators inserted at the *Sox2* locus to detect promoter-centered chromatin contacts at high resolution. In control mES cell clones with no insertion, contact frequencies between the *Sox2* promoter and downstream *SE* were similar between the CAST and 129 alleles (Fig. 4a). Inserting two CBSs from the *Sox9*-*Kcnj2* TAD boundary between the *Sox2* promoter and *SE* reduced the promoter-enhancer contacts significantly (Fisher exact test, *P* = 4.91e-4) (Fig.4a). Consistent with the observed dosage-dependent insulation effects, the *Sox2* enhancer-promoter contacts on the CAST allele were further reduced in cells with the insertion of four CBSs (Fisher exact test, *P* = 5.34e-5) (Fig. 4a). By contrast, placing two or four CBSs downstream of the *Sox2* enhancer did not reduce the *Sox2* enhancer-promoter contacts (Fig. 4a). These results support the model that insulators act by reducing the enhancer-promoter contacts.

**Fig. 4.**
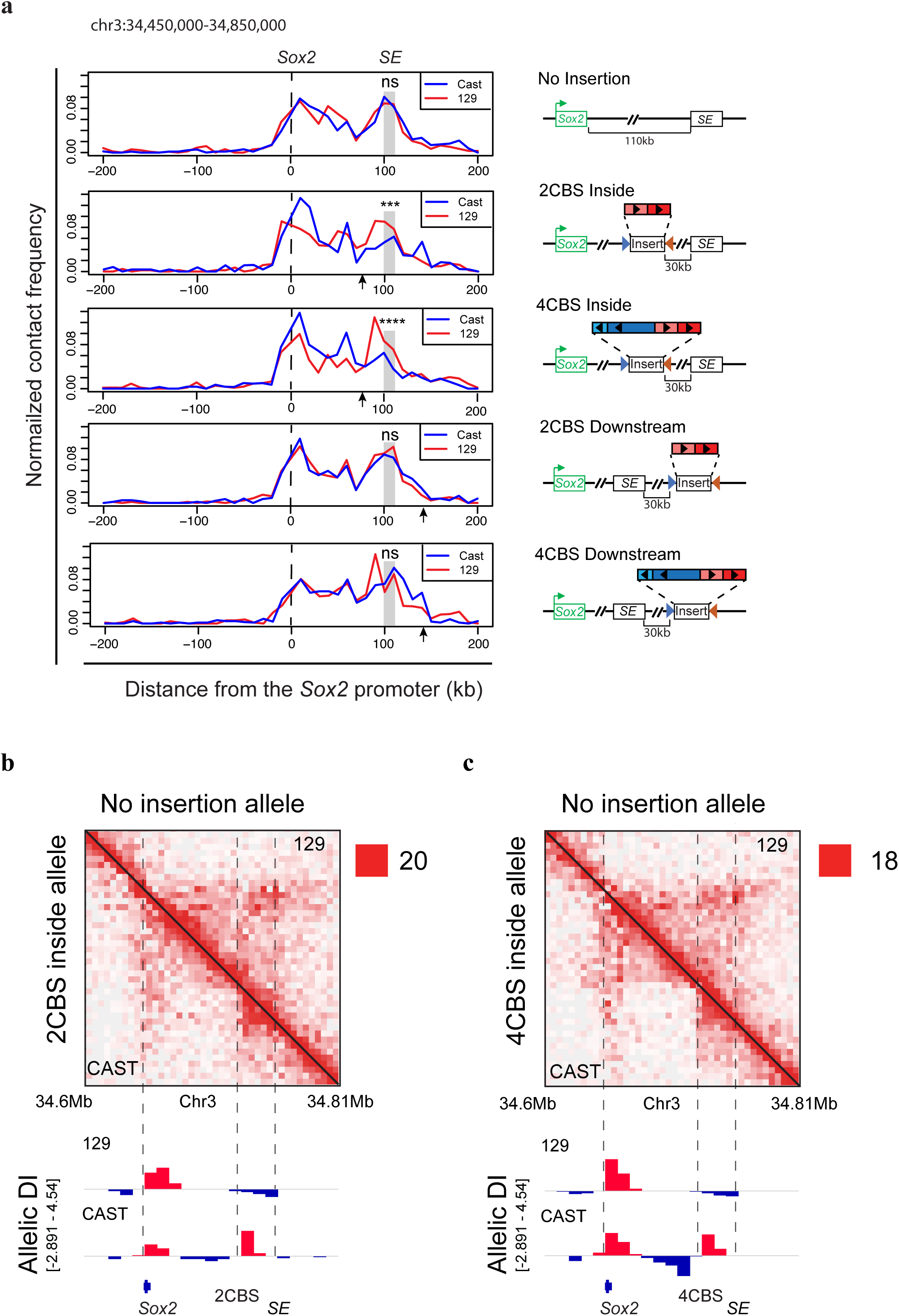
Enhancer-blocking insulator forms local chromatin domains and reduces *Sox2* enhancer-promoter chromatin contacts. **a**, Allelic chromatin contacts from PLAC-seq data are shown at the viewpoint of the *Sox2* promoter (n=2, replicates were merged). PLAC-seq reads were mapped to the mm10 reference genome and split to CAST and 129 allele based on the haplotypes of parental strains. Ambiguously mapped reads were discarded. Interaction frequency was normalized by total *cis*-contacts of the *Sox2* promoter for each allele, bin size = 10kb. Arrows indicate insertion location of CBSs. Fisher exact tests of *Sox2* enhancer-promoter contacts of the two alleles were performed. ns *P* > 0.05, \*\*\**P* = 4.91e-4, \*\*\*\**P* = 5.34e-5. Right, insertion construct matching each clone on the left. The CBS clusters were obtained from the *Sox9*-*Kcnj2* TAD boundary by PCR. **b-c**, Allelic Hi-C contact map at *Sox2* locus. Mouse ES cells with insertion of two CBSs or four CBSs from the *Sox9-Kcnj2* TAD boundary in the CAST allele were used for the experiments. Hi-C reads were mapped to the mm10 reference genome and split to CAST and 129 allele based on the haplotypes of parental strains. Ambiguously mapped reads were discarded. Allele-specific contact matrix was normalized by K-R matrix balancing. Top right, no insertion allele (129); Bottom left, insertion allele from the same cells (CAST). Bottom, allelic directionality index (DI) score of Hi-C interaction frequency (n=2, replicates were merged).

To further understand the effect of the inserted insulators on local chromatin structure, we performed *in situ* Hi-C experiments^56^ with mES cell clones containing either two or four CBSs inserted between the *Sox2* gene and its *SE* on the CAST allele (Fig. 4b-c). On the 129 allele, *Sox2* promoter and downstream *SE* were found to be in a single TAD and characterized by strong local chromatin contacts (Fig. 4b). By contrast, the insertion of two CBSs between the *Sox2* gene and *SE* on the CAST allele created a new TAD boundary that separated the *Sox2* locus into two local chromatin domains, evidenced by a sharp transition of the Directionality Index (DI) at the insertion site (Fig. 4b). Introducing four CBSs in the same location created an even stronger TAD boundary, as the transition of DI was more drastic and contacts across the new local domains were further reduced (Fig. 4c). Collectively, these results suggest that insulators create a local domain boundary between promoter and enhancer sites.

### Direct visualization of insulator-mediated changes of chromatin topology by multiplexed DNA FISH

To directly visualize the impacts of insulators on chromatin architecture, we used the recently developed multiplexed DNA FISH imaging method to trace the chromatin conformation, which allowed for visualization of the 3D organization of chromatin in single cells at tens of nanometer resolution^57-59^. We traced the 3D structure of the 210-kb genomic region (chr3: 34601078-34811078) containing the *Sox2* and *SE* loci across thousands of individual chromosomes at 5-kb intervals. We partitioned the 210-kb region into forty-two 5-kb segments and designed a library of primary oligonucleotide probes, each containing a target sequence for hybridizing to one of the 42 segments and a readout sequence that is unique to each of the segments (Supplementary Tables 4 and 5). We then sequentially labeled and imaged the 42 segments in each chromosome, using 14 rounds of hybridization of readout probes with a three-color imaging scheme (Fig. 5a). The identity of the CAST allele was determined within each nucleus based on the presence of FISH signal corresponding to the 7.5-kb insulator sequence inserted into the CAST allele that was absent in the 129 allele (Fig.5a, Extended Data Fig. 6a).

**Fig. 5.**
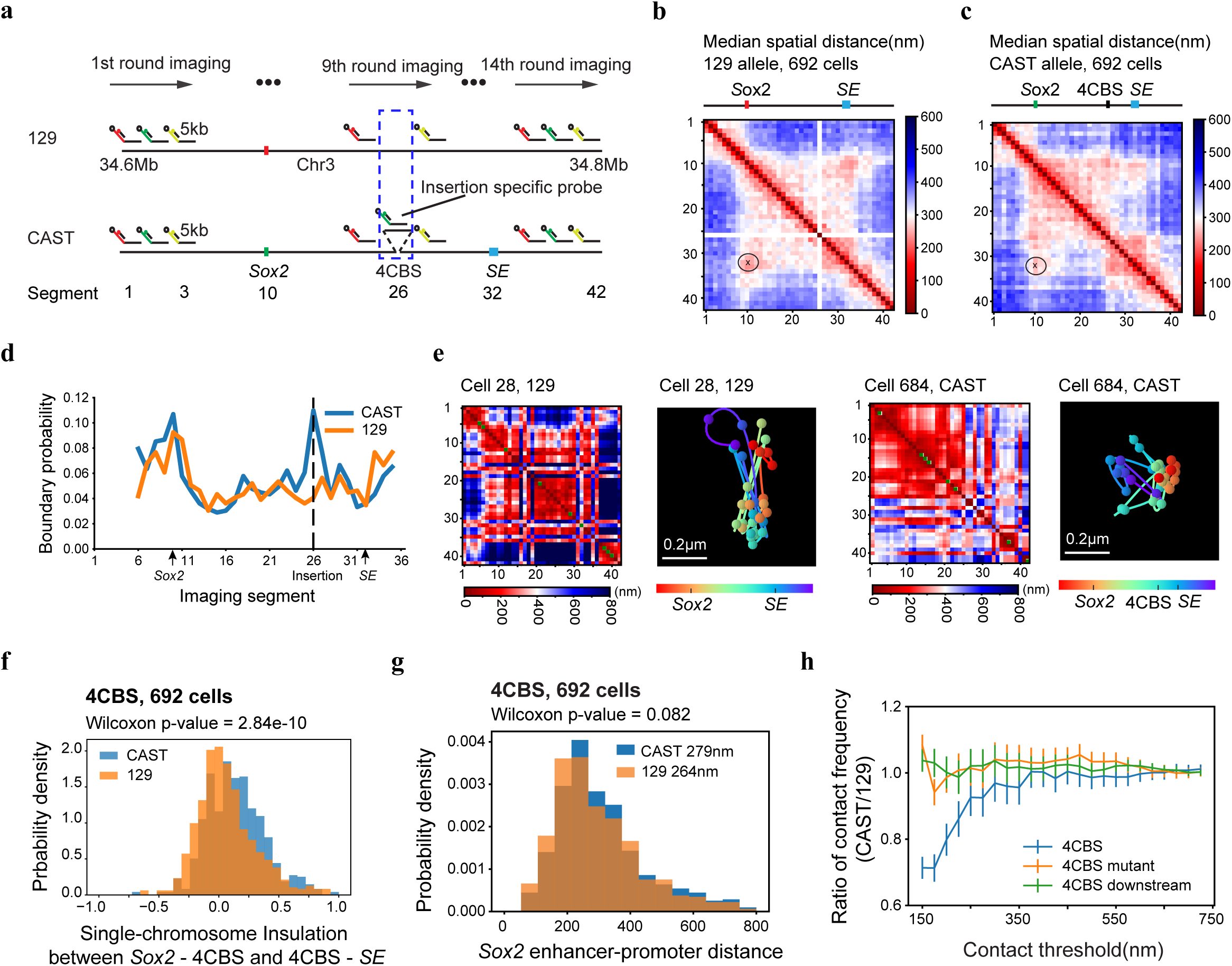
Analysis of the effects of an enhancer-blocking insulator on chromatin topology by multiplexed DNA FISH. **a**, Scheme of the chromatin tracing experiments targeting the 210-kb *Sox2* region (chr3: 34601078-34811078). Primary FISH probes were first hybridized to the entire Sox2 region. These probes were designed such that each set of probes targeting a 5-kb segment has unique readout sequences. Fluorescent readout probes were sequentially added to bind the readout sequences of each 5-kb segment via intermediate adaptor probes. Three consecutive 5-kb segments were simultaneously imaged after each round of hybridization using three color channels. 129 and CAST chromosomes in the same cell were classified based on the fluorescence signal from the insertion specific probe. The scheme shows an example of the mouse ES cell line with the insertion of 4CBS from the *Sox9-Kcnj2* TAD boundary between the *Sox2* gene and its *SE* on the CAST allele. **b-c**, Median spatial-distance matrix for the 210-kb *Sox2* region of 129 (**b**) and CAST (**c**) chromosomes from 692 cells. The 4CBS cluster was inserted between *Sox2* and its super-enhancer on the CAST allele. The 26^th^ segment was imaged by probes specific for the 4CBS insertion, therefore, it is absent from the distance matrix of the 129 alleles. **d**, The probability of each segment to be a single-chromosome domain boundary for the two alleles in **b-c**. The 26^th^ segment on the CAST allele is the 4CBS insertion. **e**, Exemplary single-chromosome structures of the imaged *Sox2* locus of CAST and 129 alleles. Interpolated single-chromosome spatial distance matrix and the matched reconstructed 3D structure are shown for each of the two alleles. Green pixels on the diagonal of the interpolated matrices indicate segments not detected in the displayed examples of chromatin traces. **f**, The distribution of single-chromosome insulation scores for each of the alleles between two domains spanning the *Sox2* promoter – 4CBS insertion (segments 10-25) and 4CBS insertion – *Sox2* enhancer (segments 26-33) regions, respectively. Insulation score was calculated for each chromosome as the natural log of the ratio of median distance between loci across domains and median distance between loci within domains. **g**, The distribution of *Sox2* enhancer-promoter distance for the CAST and 129 chromosomes in **b-c. h**, The ratio of *Sox2* enhancer-promoter contact frequency of CAST chromosomes to that of 129 chromosomes at different distance cutoffs. Contact frequency was defined as the fraction of chromosomes with *Sox2* enhancer-promoter distance below the threshold. The threshold ranges from 150nm to 750nm with 25nm intervals. The distribution of contact frequency ratio (CAST/129) of the “4CBS” clone is significantly different from that of the “4CBS mutant” and “4CBS downstream” clone, with p-value of Kolmogorov–Smirnov test equals to 6.34e-5 and 2.28e-6, respectively. The error bar represents the 95% confidence interval based upon binomial distribution.

We first carried out chromatin tracing experiments with the mES cell clone containing an insertion of the 4CBS insulator between the *Sox2* gene and the downstream super-enhancer on the CAST allele. We obtained chromatin tracing data from 692 cells where both CAST and 129 alleles were robustly discerned (Extended Data Fig. 6b, **Methods**). We then measured the spatial distance between each pair of the 5-kb genomic segments, determined the median distances across all individual chromosomes in these cells, and constructed a median spatial distance matrix for all segment pairs. Consistent with results from Hi-C (Fig. 4c), the median spatial distance matrix for the 129 allele showed a single TAD harboring both the *Sox2* and *SE* loci, whereas the spatial distance matrix for the CAST allele showed two TADs with a new boundary formed at the insertion site separating the *Sox2* and *SE* loci (Fig. 5b-c; Extended Data Fig. 7a-c). Accordingly, individual CAST chromosomes were more likely to form a boundary at the 4CBS insertion (Fig. 5d-e). Moreover, the level of insulation between the two sub-regions to either side of the inserted 4CBS, containing the *Sox2* promoter and the super-enhancer was statistically significantly enhanced on the CAST alleles (Fig. 5f). Consistently, the distances between regions across the insulator were increased on the CAST allele compared to the 129 allele (Extended Data Fig. 8a).

As controls, we also performed chromatin tracing experiments with two additional mES cell lines. One of the cell lines contained the same insulator sequence as above but had all CTCF binding motifs removed. The second control cell line had the same insulator sequence inserted at an equal distance further downstream of the *Sox2* super-enhancer. We obtained chromosome tracing data on both CAST and 129 alleles from 790 and 839 cells of the two cell lines, respectively (Extended Data Fig. 6c-d). Based on FACS analyses, neither control insert reduced *Sox2* expression on the CAST allele (Extended Data Fig. 7d). Consistently, no local chromatin domain boundary was visible between the *Sox2* and *SE* loci, and spatial insulation between the *Sox2* gene and the super-enhancer was indistinguishable between the CAST and 129 alleles (Extended Data Fig. 7e-j). Interestingly, mutant CBS inserted at the same location did not increase the distance between regions across the insertion (Extended Data Fig. 8b). In contrast, the 4CBS insulator inserted downstream of the *Sox2* super-enhancer appeared to promote segregation of the *Sox2* domain from downstream chromatin, which may explain the slightly increased Sox2 expression in this clone (Extended Data Fig. 8c).

Surprisingly, although the 4CBS insulator substantially reduced *Sox2* expression and the contact frequency between *Sox2* and its super-enhancer, the median spatial distance between *Sox2 SE* and promoter only mildly increased on the CAST alleles (279nm) compared to the 129 alleles (264nm) (Wilcoxon rank sum test, *P* = 0.082) (Fig. 5g). We hypothesized that only on a small fraction of chromosomes the *Sox2* super-enhancer was in physical proximity with the *Sox2* promoter to engage in productive transcription, and insertion of an insulator on the CAST allele could reduce this fraction of engaged *Sox2* enhancer-promoter configuration selectively on the CAST allele. To test this hypothesis, we quantified the fraction of CAST alleles that showed a spatial distance between the *Sox2* promoter and the *SE* shorter than a particular threshold and compared to that of the 129 alleles in the same cells. Indeed, in the mES cells where the 4CBS insulator was inserted between the *Sox2* gene and *SE* on the CAST allele, the ratio between the fraction of CAST alleles with spatially proximal enhancer-promoter pairs and the fraction of 129 alleles with spatially proximal enhancer-promoter pairs was much smaller than 1, at a spatial distance threshold of 150nm, and the ratio increased gradually to 1 at a spatial distance threshold of ∼300nm (Fig. 5h). By contrast, no reduction of this ratio was observed at shorter spatial threshold in mES cell clones where CTCF motifs were deleted from the insulator, or when the insulator sequence was inserted downstream of the *Sox2* super-enhancer(Fig. 5h).

Taken together, these results support the model that insulators function by establishing local chromatin domain boundaries and reducing the frequency of productive enhancer-promoter contacts, thus modulating transcriptional activity.

## Discussion

The sequence-specific DNA binding protein CTCF plays a role in both chromatin organization and transcriptional insulation, but exactly how chromatin topology is related to transcriptional insulation remains to be understood. In this study, we developed an experimental system in the mouse embryonic stem cells to quantify the enhancer-blocking activity of insulators in the native chromatin context at the *Sox2* locus. We determined the insulator activity of a number of CTCF binding sites either alone or in various combinations, and demonstrated that potent insulation was rendered by two or more CTCF binding sites concatenated together. Importantly, we found that CTCF binding alone was insufficient to confer insulation activity, rather, sequences immediately adjacent to CTCF binding motifs were required for potent insulator function. Consistent with this observation, CTCF binding sites within TAD boundaries are more likely to function as insulators than those not located at TAD boundaries, regardless of the strength of their binding by CTCF. Finally, using two orthogonal approaches to profile chromatin architecture, we showed that CTCF likely mediates transcriptional insulation by creating local chromatin domain boundaries and reducing the frequency of productive enhancer-promoter contacts. Our results therefore provide a mechanistic insight into the link between formation of chromatin domains and CTCF mediated transcriptional insulation.

We demonstrated that several factors may be involved in CTCF-mediated transcriptional insulation in mammalian cells. First, most single CBSs showed stronger insulation effects in forward than in reverse orientation, although there was one exception to this trend. Further investigation will be necessary to determine the molecular basis for the observed biases. Second and more importantly, we found that potent insulator activity depends on additive or synergistic activities from multiple CBSs. These results implicate a different working mechanism from the *Drosophila gypsy* insulator, which was ineffective in blocking enhancer activity when two tandem copies were combined^60, 61^. However, high multiplicity of CTCF binding sites is not the only requirement for strong insulation. We found that adding nine more non-boundary CBSs to a synthetic six-CBS cluster that was ineffective in insulation was unable to bring strong enhancer-blocking activity. Through sequence swapping experiments, we showed that sequences immediately adjacent to CTCF binding motifs were necessary for enhancer-blocking function. Our results suggest that CTCF sites in the genome are not all equivalent to each other, and the dependency of CTCF-mediated insulation on both dosage and flanking sequence may explain inconsistencies in insulator activities tested in previous experiments^62^.

What factors, in addition to CTCF, may contribute to transcriptional insulation by CTCF binding sites at TAD boundaries? Recent experiments showed that the cohesin complex, which establishes chromatin loops through a loop-extrusion process, could be acetylated by ESCO1 at the CTCF binding sites that anchor long-range chromatin loops^39^. ESCO1-mediated acetylation enhances the chromatin residence time of the cohesin complex, by antagonizing WAPL-mediated unloading of cohesin from chromatin. CTCF depletion is shown to reduce the cohesin acetylation and residence time on chromatin. We speculate that the dosage of CTCF and additional factors binding to CTCF-adjacent sequences may contribute to the ESCO1-dependent acetylation of cohesin complex, thereby regulating the ability of cohesin to form long range chromatin loops and TADs on chromatin.

Our study also relates the chromatin structure involving enhancer-promoter interactions, as revealed by various 3C-based and microscopy-based experiments, to enhancer-dependent transcription. From both the 3C and imaging experiments, we found that the insertion of multiple CBS sites in tandem, with the appropriate flanking sequences, induced the formation of a TAD boundary at the insertion site and resulted in physical segregation of the enhancer and promoter. The chromatin tracing results, providing direct single-cell measurements of physical distances within the *Sox2* locus, further allowed us to characterize the structural changes induced by the inserted insulators at a variety of length scales. Our analysis supports the model that enhancers occasionally come into close proximity with target promoters to facilitate transcription and that insulator sequences can substantially reduce the frequency of productive enhancer-promoter interactions that are likely within 300nm distance.

## Supporting information

SupplementaryTable1_CBS_cloned

SupplementaryTable2_Synthetic_CBS

SupplementaryTable3_tandemly_arrayed_CBSs

SupplementaryTable4_mSox2_probelibrary

SupplementaryTable5_probes_readout

SupplementaryTable6_normalizedSOX2_EXP

SupplementaryTable7_PCR_primers

## Author contributions

This study was conceived by B.R, H.H. B.R supervised the study. H.H performed insulator assays and related analysis. R.H and M.Y performed PLAC-seq/HiChIP and Hi-C experiments. I.J and M.H analyzed PLAC-seq. Y.Z performed Hi-C analysis. Q.Z and Y.H performed chromatin tracing experiments with help from B.B and X.Z. A.P.J, B.B, C.K, M.C, S.B, A.C, M.N, analyzed chromatin tracing data. The manuscript was written by H.H, B.R with input from all co-authors.

## The authors declare

Bing Ren is a co-founder and consultant for Arima Genomics, Inc. Xiaowei Zhuang is a co-founder and consultant for Vizgen, Inc.

## Acknowledgements

We are grateful for comments from members of the Ren laboratory. This study was supported by funding from the Ludwig Institute for Cancer Research and NIH (U54 DK107977, to B.R, M.H, and M.N, and 3U54DK107977-05S1 to B.R.). X.Z is a Howard Hughes Medical Institute Investigator.

## Methods

### Cell culture

The hybrid F123 mES cell line (F1 *Mus musculus castaneus* × S129/SvJae, maternal 129/Sv, paternal CAST) was from Dr. Rudolf Jaenisch’s lab at the Whitehead Institute at MIT. The wild type F123 mES cell line and engineered clones were maintained in feeder-free, serum-free 2i conditions (1uM PD03259010, 3uM CHIR99021, 2mM glutamine, 0.15uM Monothioglycerol, 1000U/ml LIF). The growth medium was changed every day. Cells were dissociated by Accutase (AT104) and passaged onto 0.2% gelatin-coated plates every 2-3 days.

### Genetic engineering of the *Sox2* locus

Tagging of the *Sox2* gene with fluorescence reporter was performed by CRISPR-Cas9 mediated homologous recombination. Specifically, a guide RNA expression plasmid (pX330, addgene #42230) targeting the 3’ of the *Sox2* gene, together with *egfp* and *mCherry* donor plasmids were co-electroporated into wild-type F123 cells by Neon transfection system (MPK1096). Cells were recovered for 2 days, then eGFP^+^ mCherry^+^ cells were sorted by FACS and seeded onto a new 0.2% gelatin-coated 60mm dish. 5 days later, a second round of FACS was performed to enrich eGFP^+^ mCherry^+^ cells.

500-1,000 double positive single cells were seeded onto a new 60mm dish and single colonies were picked manually another 5 days later. Allele-specific genotyping of *Sox2* was performed with primers spanning CAST/129 SNPs.

mCherry_Forward: CGTGGAACAGTACGAACGCG

egfp_Forward: GTCCTGCTGGAGTTCGTGAC

Reverse (common): AGAACGCTCGGCGCGTCTACTT

A clone with the CAST allele *Sox2* gene fused with *egfp* and 129 allele *Sox2* gene fused with *mCherry* was selected as the parental clone. Subsequently, the *HyTK* fusion gene was integrated into the CAST allele of the parental clone by CRISPR-Cas9 editing. Specifically, electroporated cells were recovered for 2 days and then cultured in growth media containing 200ug/ml hygromycin for 7 days. Survived cells were dissociated into single cells and seeded at the density of 500-1,000 cells per 60mm dish. 5 days later, colonies were manually picked and genotyped with primers spanning CAST/129 SNPs. Genotyping primers of *HyTK* fusion gene for insulator reporter and control cell lines: Inside_F: GGAGCTCACCGATTATGTGC

Inside_R: GAACTTCGGATCCACTGAAAACA

Downstream_F: GGATGGTCCAGACCCACGTC

Downstream_R: AGATGCTCTGTCGGTCACTG

### Donor plasmids cloning for recombinase mediated cassette exchange (RMCE)

The donor vector was adapted from the pUC19 plasmid. Two heterotypic Flippase recognition sites FRT(GAAGTTCCTATTCCGAAGTTCCTATTCTCTAGAAAGTATAGGAACTTC), F3(GAAGTTCCTATACTATTTGAAGAATAGGAACTTCGGAATAGGAACTTC), as well as NotI and SbfI restriction enzyme recognition sites, were added into pUC19 plasmid by PCR. The donor vector was then digested with the enzyme cocktail of NotI-HF (neb, R3642S), SbfI-HF(neb, R3189S), and rSAP(neb, M0371S) for 4hrs at 37 °C. Individual CTCF binding sites were PCR amplified from mouse or human genomic DNA. PCR primers contain overhang sequences of NotI and SbfI sites to specify CTCF motif orientation. PCR products were purified by gel-electrophoresis and digested by NotI-HF and SbfI-HF at 37°C for 30min. The digestion mix was then inactivated at 65 °C for 20min, purified with SPRI beads (1:1 ratio) and ligated into the digested donor vector. Ligation products were transformed into Stbl3 chemically competent cells. Positive clones were screened by PCR and inoculated in 50ml of LB at 37 °C for 16 hours. Plasmids were extracted using QIAGEN plasmid plus midi kit (cat 12943) and validated by sanger sequencing.

### Genetic engineering of insulator reporter mESC by RMCE

A Flippase expression plasmid(pFlpe) (addgene #13787) and a donor plasmid(pDonor) were co-electroporated into 0.1 million insulator reporter or control cells at the ratio of 1:4 (pFlpe: pDonor = 1μg :4μg). Cells were seeded onto a 6-well plate and recovered for two days. Then, cells were cultured in growth media containing 2μM ganciclovir for 5 days. Survived cells were dissociated into single cell suspension and seeded at the density of 500-1,000 cells per 60mm dish. Five days later, six colonies were picked for PCR genotyping. Genomic DNA was then extracted by QIAGEN DNeasy Blood & Tissue Kits (#69506, #69581). For each insert, three independent clones were randomly picked for FACS analysis and subsequent studies.

Genotyping primers for insertion in insulator reporter and control cell lines:

Inside_F: GGAGACAAGAGATGTCAGGAG

Inside_R: TCCGCAAGCAAATAGCTCCATTC

Downstream_F: CATCGGCAATGAGTGTGTGTCA

Downstream_R: GTGATCTCCAGAGTATACGCATGTC

Individual CTCF binding sites were combined by PCR to create CBS clusters. Specifically, the 4CBS cluster from the *Sox9-Kcnj2* TAD boundary was consisted of genomic sequences from chr11:111,523,291-111,524,273, chr11:111,531,104-111,533,964, and chr11:111,535,307-111,538,959.

### FACS data acquisition and analysis

Cells were treated by Accutase(#AT104) at 37°C for 5-7min and resuspended into single cells with 2ml warm 2i/LIF medium. Cells were then spun down at 1,000rpm for 4min and washed twice with 5ml PBS. Cell pellets were resuspended into single cells with 1ml PBS and filtered through the 35μm strainer cap of a FACS tube (SKU: FSC-9005). Then, cells were sorted by Sony sorter SH800 in analysis mode using a 130μm chip. For each insertion clone, both GFP and mCherry signals were recorded for 10,000 cells. Multiple technical replicates of the no insertion clone was included as controls for every FACS sorting experiment. Cells were first gated by SSCA-FSCA for live cells, then by FSA-FSH for singlets. Fluorescence signals of cells passed gating were exported in csv files and analyzed in R. Specifically, the GFP signal is normalized by mCherry signal from the same cell. For each insertion clone, the normalized Sox2-eGFP expression was calculated as:

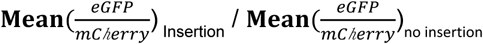

To better estimate instrument variability in FACS sorting, we used replicates of the no insertion clone in all experiments as controls when testing the significance of insulation effects of the inserted DNA elements.

### ChIP-seq

ChIP-seq was performed as previously described with minor modifications^63^. Briefly, cells were dissociated into single cells and cross-linked by 1% formaldehyde in PBS for 15min at room temperature. Cross-linking was then quenched by 0.125M glycine and cells were washed twice with 5ml cold PBS. Permeabilized nuclei were prepared with Covaris truChIP Chromatin Shearing Kit (PN520154) following the manufacturer’s instructions. 1-3 million nuclei were sonicated in 130μl microtube by Covaris M220 instrument (Power, 75W; Duty factor, 10%; Cycle per bust, 200; Time, 10 mins; Temperature, 7°C.). Sonicated chromatin was diluted with 1xShearing Buffer into a total volume of 1ml and spun down at 15,000rmp at 4°C to remove cell debris. 5ug antibodies were added to the supernatant and incubated overnight at 4°C with gentle rotation (CTCF, ab70303; RAD21, ab992; H3K4me3, Millipore, 04-745; H3K27ac, Active Motif,39685.). Chromatin was pulled down by protein G Sepharose beads (#17061801, GE health care) and washed three times with RIPA buffer(10 mM Tris pH 8.0, 1 mM EDTA, 1% Triton X-100, 0.1% SDS, 0.1% Sodium Deoxycholate), two times with high-salt RIPA buffer (10 mM Tris pH 8.0, 300 mM NaCl, 1 mM EDTA, 1% Triton X-100, 0.1% SDS, 0.1% Sodium Deoxycholate), once with LiCl buffer (10 mM Tris pH 8.0, 250 mM LiCl, 1 mM EDTA, 0.5% IGEPAL CA-630, 0.1% Sodium Deoxycholate), and twice with TE buffer (10 mM Tris, pH 8.0; 0.1 mM EDTA). Washed chromatin was reverse crosslinked overnight with 2μl proteinase K (P8107S, NEB) at 65 °C (1%SDS, 10 mM Tris, pH 8.0, 0.1 mM EDTA). Reverse-crosslinked DNA was column purified and subjected to end repair, A-tailing, adapter ligation, and PCR amplification. Final libraries were purified by SPRI beads (0.8:1) and quantified with Qubit HS dsDNA kit (Q32854) prior to Illumina next-generation sequencing.

### PLAC-seq/HiChIP

Proximity Ligation ChIP-sequencing (PLAC-seq) (also known as HiChIP) libraries were prepared as previously described^54, 55^ with minor modifications. In brief, 2-3 million cells were crosslinked for 15 minutes at room temperature with 1% methanol-free formaldehyde and quenched for 5 minutes at room temperature with 0.2 M glycine. The crosslinked cells were lysed in 300 μl Hi-C lysis buffer (10 mM Tris-HCl, pH 8.0, 10 mM NaCl, 0.2% IPEGAL CA-630) for 15 minutes on ice and then washed once with 500 μL lysis buffer (2,500xg for 5 minutes). Subsequently, cells were resuspended in 50 μl 0.5% SDS and incubated for 10 mins at 62°C then quenched by 160 μl 1.56% Triton X-100 for 15 mins at 37°C. Then, 25 μl of 10X NEBuffer 2 and 100 U MboI were added to digest chromatin for 2 hours at 37°C with shaking (1,000 rpm). Enzymes were inactivated by heating for 20 mins at 62°C. Digested fragments were biotin-labeled and subsequently ligated by T4 DNA ligase buffer (NEB) for 2 hours at 23°C with 300 rpm gentle rotation. Chromatin was sheared and washed as described in ChIP-seq. Dynabeads (M-280 Sheep anti-Rabbit IgG) coated with 5μg H3K4me3 antibodies (Millipore, 04-745) were used for immunoprecipitation. Pulled down chromatin was treated with 10 μg RNase A for 1 hour at 37°C, and subsequently reverse-crosslinked by 20 μg proteinase K at 65°C for 2 hours. DNA fragments were purified with Zymo DNA Clean & Concentrator-5 kit. Ligation junctions were enriched by 25 μl myOne T1 Streptavidin Dynabeads. Libraries were prepared using QIAseq Ultralow Input Library Kit (Qiagen, #180492). Final libraries were directly PCR amplified from Streptavidin beads, size selected with SPRI beads (0.5:1 and 1:1), quantified and submitted for paired-end sequencing.

### Hi-C

Cells were processed in the same way as in PLAC-seq before chromatin shearing steps. Briefly, nuclei after the ligation step were digested by 50 μl of proteinase K (20mg/ml) for 30min at 55 °C. DNA was then purified by ethanol precipitation and resuspended in 130μL 10mM Tris-HCl (PH=8.0). Purified DNA was sonicated by Covaris M220 instrument with the following parameters: Duty cycle, 10%; Power, 50; Cycles/burst, 200; Time, 70 seconds. DNA fragments smaller than 300bp were removed by Ampure XP bead-based dual size selection (0.55:1 and 0.75:1). Biotin-labeled free DNA ends were cleaned up by end-repair reaction and ligation junctions were enriched by Streptavidin Dynabeads as described in PLAC-seq. Ligation junctions were then purified and subjected to A-tailing, adapter ligation, and PCR amplification. Final libraries were purified by 0.75x Ampure XP beads, quantified and submitted for pair-end sequencing.

### Multiplexed FISH imaging for chromatin tracing

Glass coverslips were treated by poly-L-lycine for 30min at 37°C. Then, glass coverslips were washed twice with 5ml PBS and treated by 0.2% gelatin for another 20min at 37°C. 2.5 million mouse ES cells were seeded in a 6cm plastic dish containing the treated glass coverslip. After 20 hours, cells were cross-linked by 4% paraformaldehyde and followed by chromatin tracing experiments as described in a previous publication^57^. Briefly, the entire 210kb Sox2 region was labeled by a library of primary Oligopaint probes^57, 58^. Each primary probe consists of a unique 42-nucleotide readout sequence that is specific for each 5kb DNA segment. Next, secondary readout probes complementary to the readout sequences on the primary probes were added to the cells. Lastly, fluorophore-labeled common imaging probes complementary to the secondary probes were added to the cells to allow 3D diffraction-limited imaging of individual DNA segments. After each round of imaging, the fluorescence signal was extinguished by using both TCEP [tris(2-carboxyethyl) phosphine] cleavage at a concentration of 50μM in 2x SSC and high power photobleaching. The process was repeated until all DNA segments were labeled and imaged. To increase the throughput, we performed three-color imaging by using three secondary readout imaging probes that were conjugated with Cy3, Cy5, and Alexa 750, respectively. In this case, three consecutive 5-kb chromatin segments were labeled by each round of imaging. A pool of 42 oligo probe sets was designed to scan the 210kb *Sox2* locus with each set covering a 5 kb DNA region. The 7.5kb 4 CBS insertion was imaged by the 26th probe set.

## Data analysis

### ChIP-seq

Sequenced reads were aligned to reference mouse genome mm10 and unmapped reads and PCR duplicates were removed. For clones with the insertion of synthetic CTCF binding sites, reads were aligned to a customized mm10 reference genome that includes the inserted sequence. Signal tracks were generated with the command “bamCoverage –normlizingRPKM -bs 50 --smoothLength 150”. Allele-specific reads were resolved based on SNP VCF files described in the PLAC-seq analysis below.

### PLAC-seq

To resolve allele-specific interactions, we created the VCF files containing SNPs with respect to the mm10 reference genome for parental strain CAST/EiJ and 129SV/Jae. Specifically, whole-genome sequencing reads from the two strains were mapped to mm10, deduplicated, and called SNPs using bcftools. Since parental strains are highly inbred and should be homozygous for all sites, we removed heterozygous SNP calls and those with sequencing depth less than 5 and quality less than 30. We further removed SNPs that were present in both strains. In the end, we kept 19863797 distinguishable SNP sites for the two alleles of the F123 cell line. We used a modified mapping procedure from WASP^64^ pipeline to detect allele-specific contacts. Since WASP pipeline ignores indels, we further removed all reads which map to within 50 base pairs from the nearest indel. Briefly, paired-end reads were first mapped to mm10 reference genome, and reads overlapped with polymorphism sites were remapped after changing the nucleotide at the SNP’s position to match the other allele. If such, ‘flipped’, reads were mapped to the same position as before, reads were kept and assigned to either maternal or paternal allele based on SNP information. Otherwise, the reads were discarded. For duplicated reads, instead of choosing the read with the highest mapping score, a random read was kept. We modified the original WAPS mapping procedure by replacing the bowtie2 alignment tool with bwa-mem and integrated MAPS^65^ feather post-filtering pipeline to resolve the chimeric reads.

### Hi-C

To process Hi-C data we used our in-house pipeline available at https://github.com/ren-lab/hic-pipeline. Briefly, Hi-C reads were aligned to mm10 using BWA-MEM for each read separately and then paired. For chimeric reads, only 5’ end-mapped locations were kept. Duplicated read pairs mapped to the same location were removed to leave only one unique read pair. The output bam files were transformed into juicer file format for visualization in Juicebox. Contact matrices were normalized using the Knight–Ruiz matrix balancing method^66^. Directionality Index (DI) score for each sample was generated at 50-kb resolution and 2-Mb window (40 bins) as described in a previous work^25^. Haplotype phasing was performed using the obtained Cast/129 VCF file. This created two contact matrices corresponding to ‘Cast allele’ and ‘129 allele’ for each Hi-C library. For each phased haplotype of chromosome 3, the DI score was generated at 10-kb resolution and 50-kb window (5 bins).

### Chromatin tracing data processing

Custom software was used to obtain images of chromatin architecture as described previously^57^ with minor modifications. The software identifies centroid positions of each 5-kb chromatin segment using diffraction-limited z-stack images acquired by epifluorescence microscopy. Chromosome locations were first identified via the segmentation of the nuclei (stained with DAPI) in each field of view using a convolutional neural network (CNN). The segmentation masks were then applied to limit the chromosome candidates to the two most likely clusters of fluorescence spots presented in each nucleus. We then selected the two spots that showed strongest averaged fluorescence signal over all imaging rounds as the two alleles for each nucleus. To avoid selecting the same chromosome, we also required the two spots to be separated by at least 10 pixels (1.08μm). The algorithm then utilized the identified chromosome locations to select candidate spots of the imaged 5-kb chromatin segments in every round of imaging. A Gaussian fitting algorithm was then used to fit both the signal of each of the candidate segments and the fiducial beads. The chromatic aberration, flat-field, and drift correction algorithms were adopted from the published work^57^.

To minimize misidentification of fluorescence spots, the candidate spot of each segment was then further evaluated for their likelihood to be accepted or rejected as estimated by an expectation maximization (EM) algorithm. The EM algorithm computes a score based upon a product of three terms which measure the relative rank, from 0 to 1, of each candidate spot of a segment among all candidates within the 3-D window centered upon the chromosome location. The three terms measure the brightness of the spot, the proximity of the spot to the estimated chromosome centroid position, and the proximity of the spot to a moving average localization of the candidates selected in the previous five rounds of imaging. This scoring scheme enables selection based upon a segment’s similarity to other high-quality segments. It also allows for dimmer candidate spots to be considered with confidence if the local environment is sufficiently clear of noise. The EM algorithm selected the highest scoring candidate spot for each chromosome segment in each round of imaging, while all remaining candidate spots were not considered in subsequent analyses.

With the scores computed, we then identified a threshold which resulted in a chromosome misidentification rate below 10%. The misidentification rate was computed as the percentage of fluorescence spots among the top discarded candidate spots which had scores above the EM score threshold that we chose. Finally, only chromosomes that contained accepted segments with a score above the selected threshold across at least ∼50% of imaging rounds (22/42 rounds) were kept for further analysis. The detection efficiency of each segment for each experiment was computed as the fraction of segments with accepted candidate spots based upon the above procedure, which was around 64% for all experiments. To avoid misclassification of the two alleles in the same mES cells, we only kept cells in which one and only one chromosome was detected positive for the insertion. Any nuclei showed fluorescence signal on both alleles or neither allele for the 7.5kb insertion were discarded. In this way, misclassification of the two alleles is estimated to be less than 5%. Then, pairwise distances between each 5kb segment were computed for each chromosome. The resulting matrices were combined into an aggregate distance matrix for each allele by taking the median value across all chromosomes within each group (CAST or 129). To compute the single chromosome insulation score, we employed the methods and algorithms described in previous work^57^. Sox2 enhancer-promoter distance was calculated by median pairwise Euclidean distances between the genomic locations of the *Sox2* gene (9^th^ - 11^th^ region) and its enhancer (30^th^ - 32^nd^ region) for every chromosome.

## DATA access

To review GEO accession GSE153403: Go to https://www.ncbi.nlm.nih.gov/geo/query/acc.cgi?acc=GSE153400.

## Extended figure legends

**Extended Data Figure. 1.**
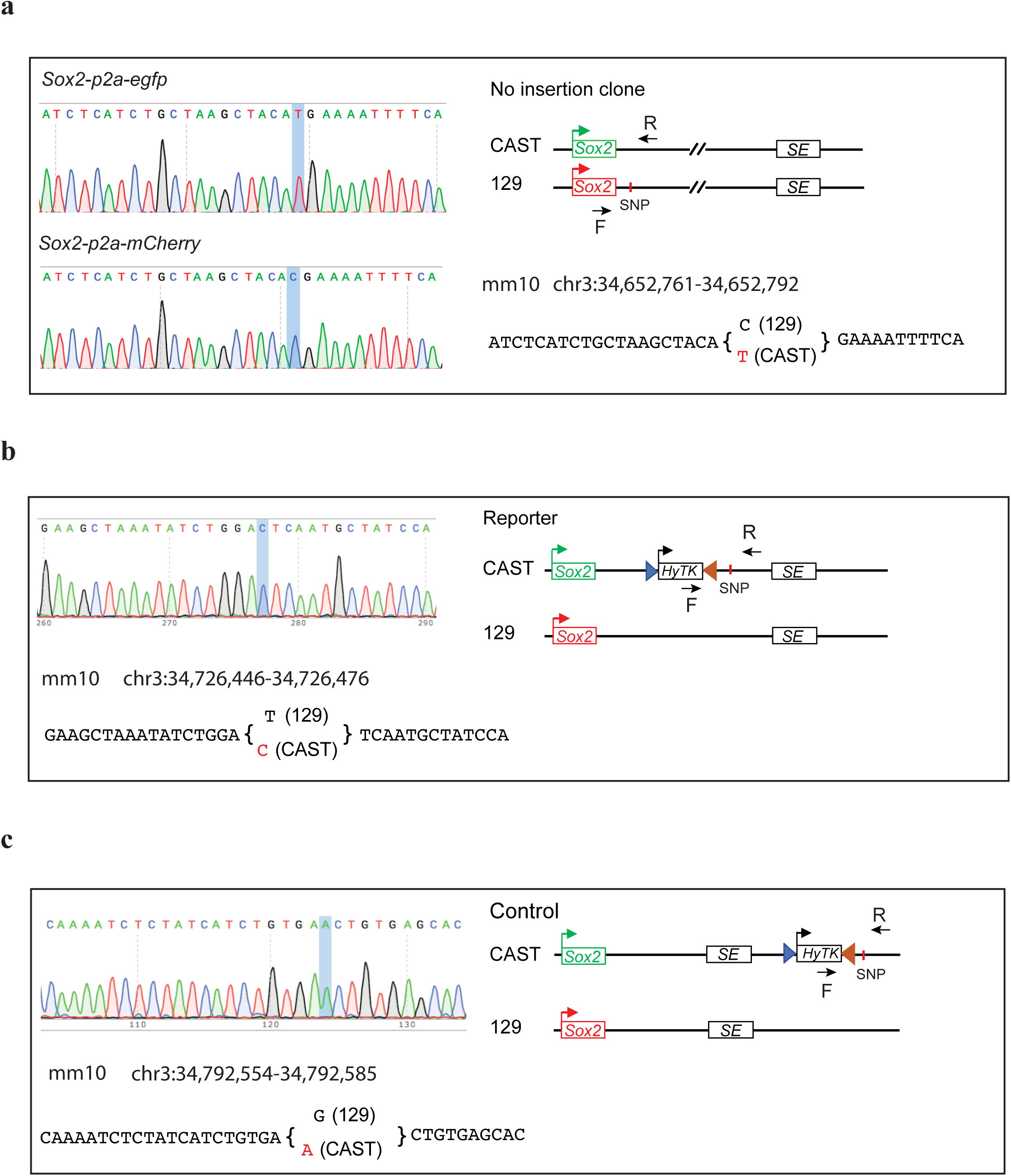
Genotyping the engineered mES cell lines. **a**, Genotyping *egfp* and *mCherry* labeled *Sox2* gene. Left, Sanger sequencing results for allele-specific PCR products. Allele-specific SNP is highlighted. Right, Construct of the clone and the SNP information used to distinguish the two alleles. The reverse primer was common, while the forward primer was allele-specific, matching with *egfp* and *mCherry* sequence, respectively. **b-c**, Genotyping the Insulator reporter and control cell lines. Left, Sanger sequencing and SNP information. Right, Construct of the clone and positions of PCR primers. The forward primer is specific to the inserted *HyTK* gene. **b**, insulator reporter cell line. **c**, Insulator control cell line.

**Extended Data Figure. 2.**
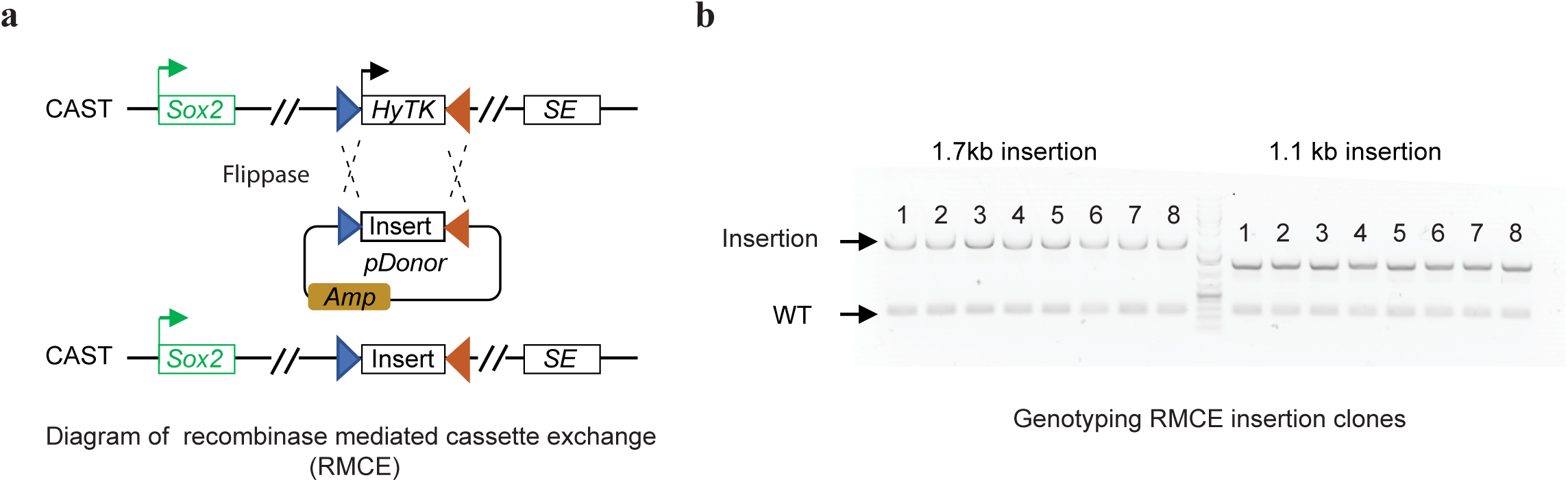
Efficiency of insertion by recombinase mediated cassette exchange. **a**, Diagram of recombinase mediated cassette exchange (RMCE) in the insulator reporter cell line. Flippase expression plasmid and the donor plasmid carrying the insertion sequence were co-electroporated into cells. The replacement only happens on the CAST allele. **b**, Genotyping insertion clones of λDNA fragments generated by RMCE. PCR primers were designed from genomic locations that spanned the insertion position. Top band, insertion fragment; Bottom band, PCR product from the no insertion allele.

**Extended Data Figure. 3.**
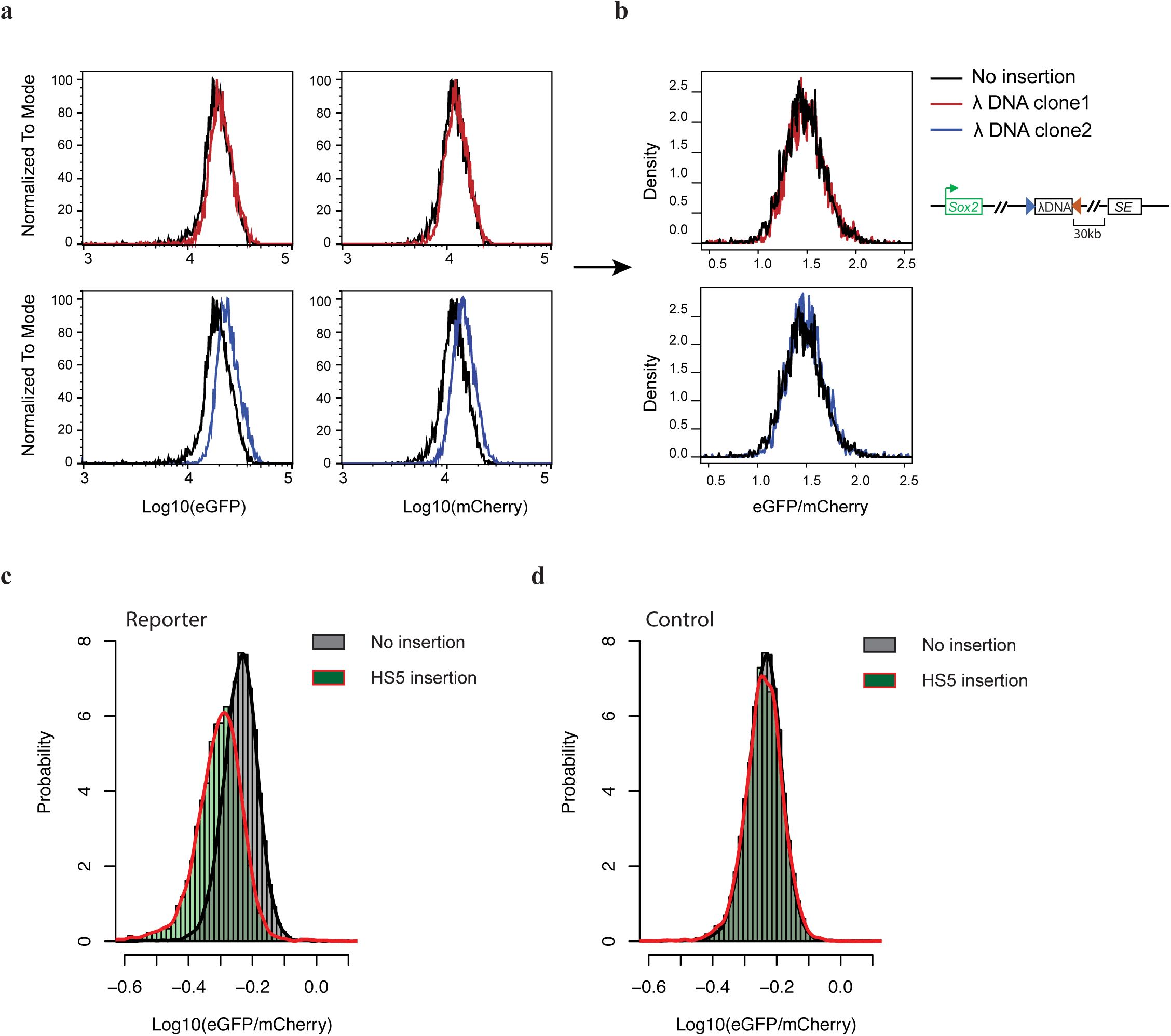
Normalization of *Sox2* expression. **a-b**. FACS profiles of two clones with the insertion of the same λDNA fragment. **a**, Histograms showing eGFP and mCherry signals of the two clones; **b**, Density plots of normalized signal (eGFP/mCherry) of cells from the two clones. For every cell, the ratio of eGFP signal over mCherry signal was calculated. **c**, A histogram shows the normalized Sox2-eGFP expression of cells with the human β-globin HS5 insulator inserted between the *Sox2* gene and its super-enhancer. The CTCF motif of the HS5 insulator was in forward orientation. **d**, A histogram shows the normalized Sox2-eGFP of cells with the human β-globin HS5 insulator inserted downstream of the *Sox2* super-enhancer. The CTCF motif of the HS5 insulator was in forward orientation.

**Extended Data Figure. 4.**
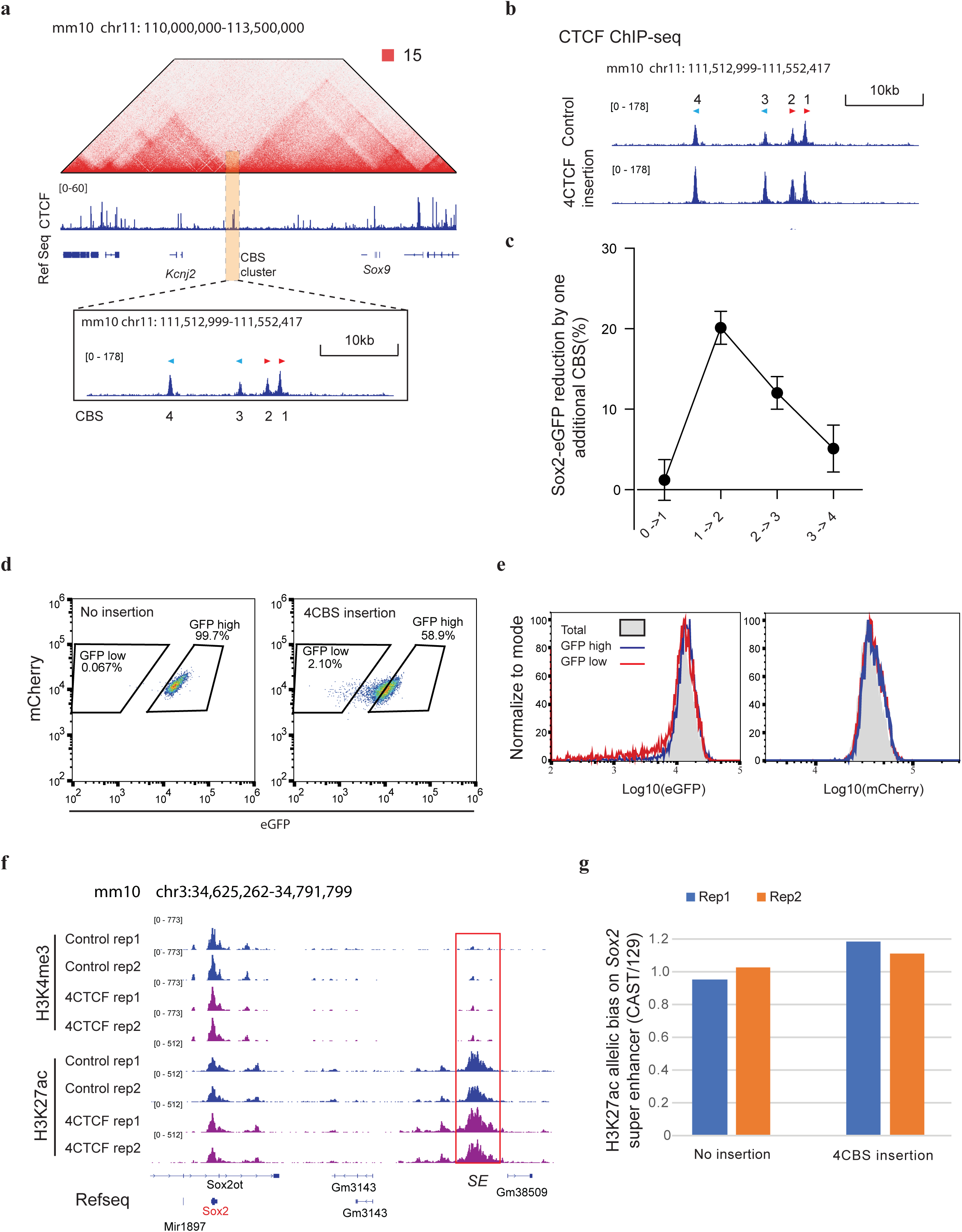
Insulation features of CBSs from the *Sox9-Kcnj2* TAD boundary. **a**, Hi-C contact map of the *Sox9-Kcnj2* locus in mouse ES cells. ChIP-seq of CTCF and RefSeq genes are shown below. CTCF binding sites at the *Sox9-Kcnj2* TAD boundary are highlighted in the orange box. Zoom in view shows the four CTCF binding sites cloned for insulator activity test. **b**, ChIP-seq of CTCF in the no insertion clone and the clone with an extra copy of the four *Sox9-Kcnj2* TAD boundary CBS inserted inside the *Sox2* domain. ChIP-seq reads were aligned to the mm10 reference genome. **c**, Reduction in Sox2-eGFP expression by one additional CBS (Data are mean ± sd). **d**, FACS profiling of the no insertion clone and the clone with the four *Sox9-Kcnj2* TAD boundary CBS (4CBS) inserted between *Sox2* and its super-enhancer. GFP^low^ and GFP^high^ sub-populations were gated. **e**, FACS profiling of GFP^low^, GFP^high^ sub-populations, and the unsort total population of the 4CBS insertion clone in **d** after extended culturing for 8 days. Left, GFP signal, right, mCherry signal from the same cells. **f**, ChIP-seq of H3K4me3 and H3K27ac in the no insertion clone and the clone with the four *Sox9-Kcnj2* TAD boundary CBS inserted inside the *Sox2* domain (n=2). The *Sox2* super-enhancer is highlighted in the red box. **g**, Allelic quantification of H3K27ac signal on the *Sox2* super-enhancer of clones in **f**. H3K27ac ChIP-seq reads on the *Sox2* super-enhancer were normalized by the total reads mapped to chromosome 3 for each allele. Then, the ratio of the normalized H3K27ac signal of the two alleles was calculated (CAST/129).

**Extended Data Figure. 5.**
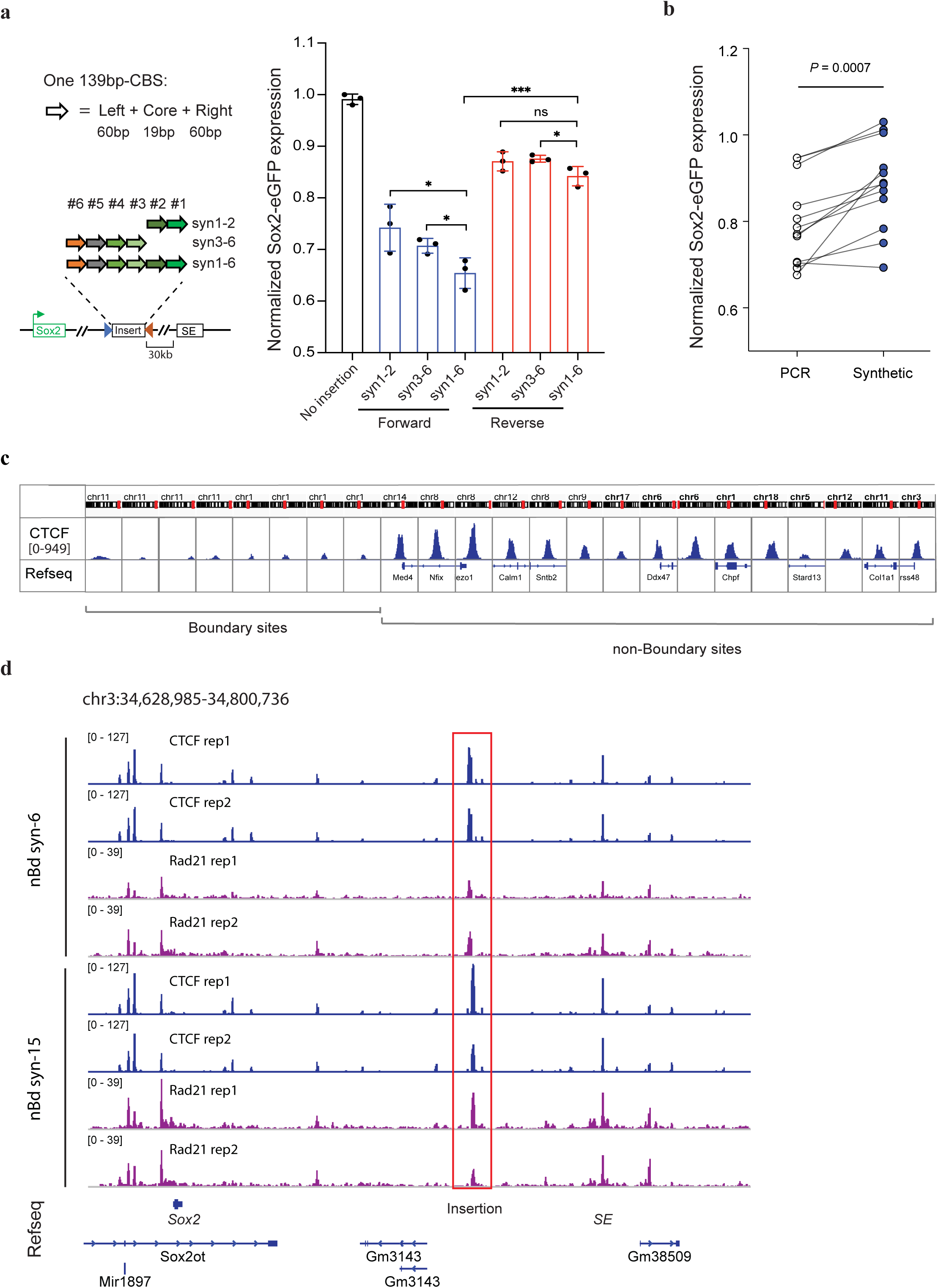
Insulation effects of synthetic CTCF binding sites. **a**, Additive insulation by synthetic CBS from boundary regions. Left top, compositions of one 139bp-CBS that was synthesized; Left bottom, tandemly arrayed 139bp-CBSs tested for insulator activity. Right, normalized Sox2-eGFP expression of clones with the tandemly arrayed 139bp-CBSs inserted between the *Sox2* gene and its super-enhancer. Blue, CBS core motifs were in forward orientation; Red, CBS core motifs were in reverse orientation. Insertions were on the CAST allele only. n=3, unpaired t-test, two-tailed. ns *P* > 0.05, \**P* ≤ 0.05, \*\**P* ≤ 0.01, \*\*\**P* ≤ 0.001, \*\*\*\**P* ≤ 0.0001. Data are mean ± sd. **b**, Insulation effects of PCR cloned large size CBSs (1-4 kb) and the synthesized 139bp-CBSs that contain the same CTCF motifs. (n=12, paired t-test, two-tailed, \*\*\**P* = 0.0007.). **c**, CTCF binding strength at selected boundary sites and non-boundary sites in mouse ES cells. ChIP-seq signals of CTCF are shown in 2-kb window. **d**, ChIP-seq of CTCF and Rad21 in clones with the insertion of six (nBd-syn6) or fifteen (nBd-syn15) 139-bp CBSs obtained from non-boundary regions. ChIP-seq reads were mapped to a customized mm10 genome that included the inserted sequence at the target site. Insertion position is highlighted in red box.

**Extended Data Figure. 6.**
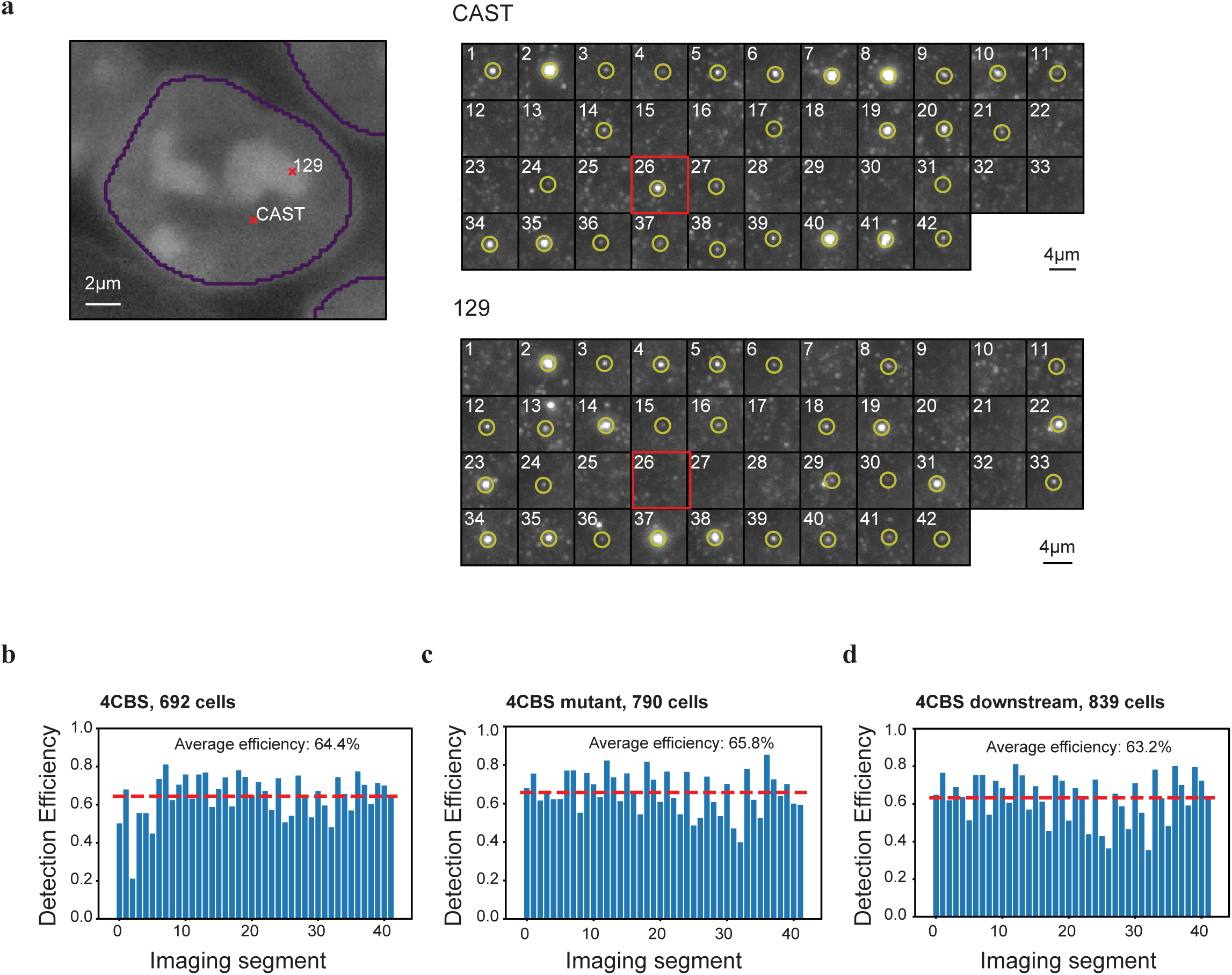
Allele classification by multiplexed DNA FISH. **a**, Exemplary images of allele classification. Left, nuclei segmentation and the positions of CAST and 129 allele in the nucleus. Right, images of the forty-two 5-kb segments (chr3:34,601,078-34,811,078) of the CAST and 129 allele. The hybridization probes of the 26^th^ segment (highlighted in the red box) specifically targeted the 4CBS sequence. The chromosome positive for the 26^th^ segment (inserted 4CBS) was classified as CAST allele, the negative chromosome in the same cell was classified as 129 allele. Cells with both chromosomes positive or both chromosomes negative for the 26^th^ segment were discarded. **b-d**, Bar plots showing detect efficiency of the 42 segments of chromatin tracing experiments in the “4CBS” clone **(c)**, the “4CBS mutant” clone **(d)**, and the “4CBS downstream” clone **(e)**. Detect efficiency of each segment was calculated as the fraction of chromosomes that showed positive fluorescence signal at the specific imaging round.

**Extended Data Figure. 7.**
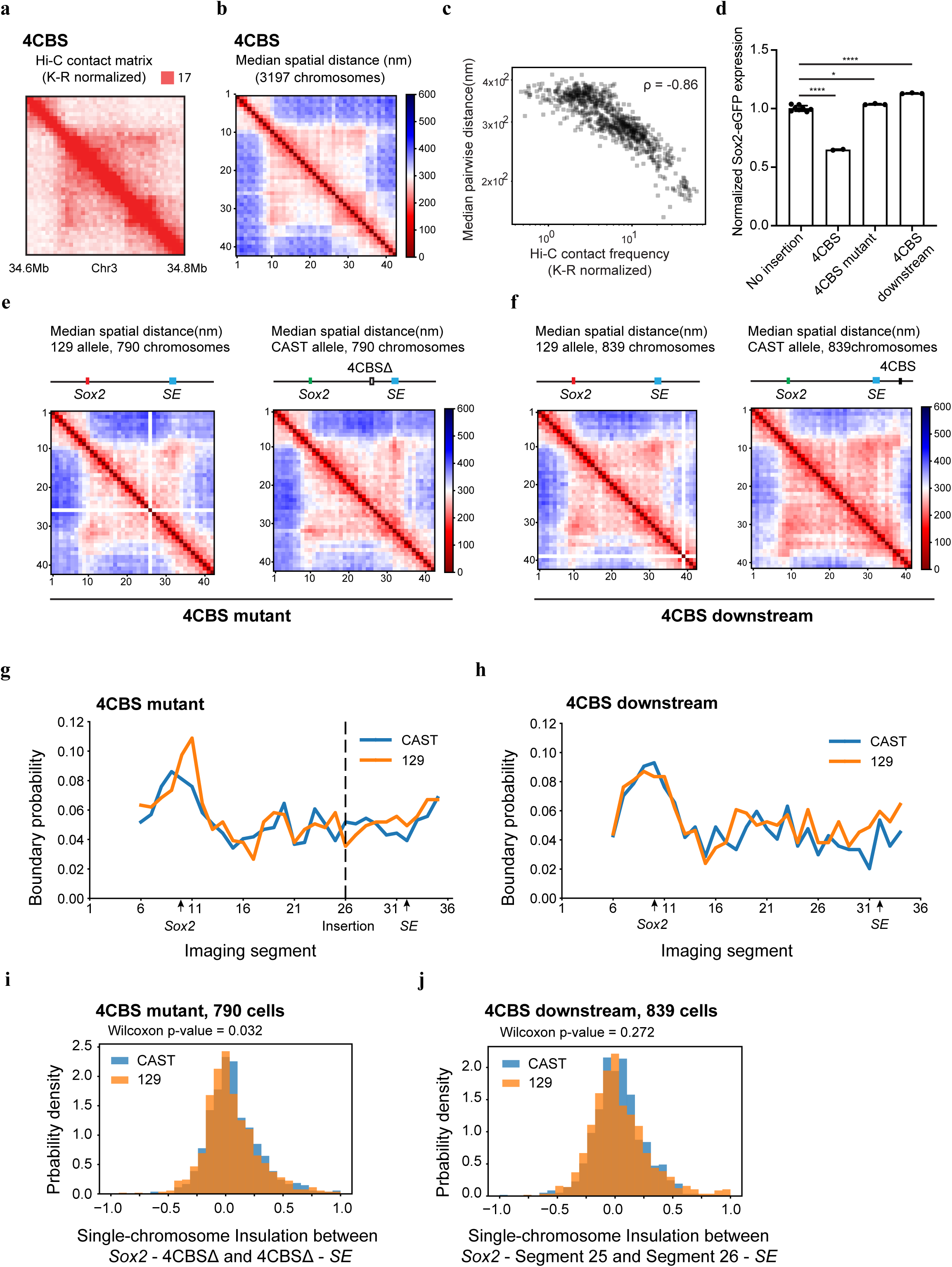
Spatial organization of the *Sox2* locus in engineered mES cells. **a**, Bulk Hi-C contact matrix (K-R normalized) of the *Sox2* locus in cells with 4CBS inserted between the *Sox2* gene and its super-enhancer on the CAST allele. **b**, Median pairwise distance of the same *Sox2* region measured by chromatin tracing experiment in the same clone in **a**, CAST and 129 chromosomes were combined. **c**, Correlation between the Hi-C contact frequency matrix (**a**) and median distance matrix(**b**). **d**, Normalized Sox2-eGFP expression in the no insertion clone(n=8), the “4CBS” clone (same cells in **a-b**, n=2), and two insertion controls. “4CBS mutant” (n=3) was the insertion clone of a 4CBS sequence that had all four 19-bp CTCF core motifs deleted (4CBSΔ). The insertion position was the same as the “4CBS” clone; “4CBS downstream” (n=3) was the insertion clone of the same 4CBS insulator sequence but located at equal distance downstream of the *Sox2* enhancer. One-way analysis of variance with Bonferroni’s multiple comparisons test. ns *P* > 0.05, \**P* ≤ 0.05, \*\**P* ≤ 0.01, \*\*\**P* ≤ 0.001, \*\*\*\**P* ≤ 0.0001. Data are mean ± sd. **e-f**, Median spatial-distance matrix for the 210kb *Sox2* region (chr3: 34601078-34811078) of 129 (left) and CAST (right) chromosomes of the “4CBS mutant” clone**(e)** and the “4CBS downstream clone” **(f)**. The 26^th^ segment was imaged by 4CBS specific probes; therefore, it is absent on the distance matrix of no insertion 129 alleles. Similarly, the 38^th^ segment is absent on the distance matrix of 129 alleles in **f. g-h**, The probability of forming single-chromosome domain boundaries at each segment for the two alleles of the “4CBS mutant” clone (**g**), and the “4CBS downstream” clone (**h**). **i**, The distribution of single-chromosome insulation scores for each of the alleles between two domains spanning the *Sox2* promoter – 4CBSΔ insertion (segments 10-25) and 4CBSΔ insertion – *Sox2* enhancer (segments 26-33) regions, respectively. Insulation score was calculated for each chromosome as the natural log of the ratio of median distance between loci across domains and median distance between loci within domains. **j**, The distribution of single-chromosome insulation scores for each of the alleles between the same two domains (segment 10-25 and segment 26-33) in (**i**) for the “4CBS downstream” clone. Insulation score was calculated in the same way as in (**i**).

**Extended Data Figure. 8.**
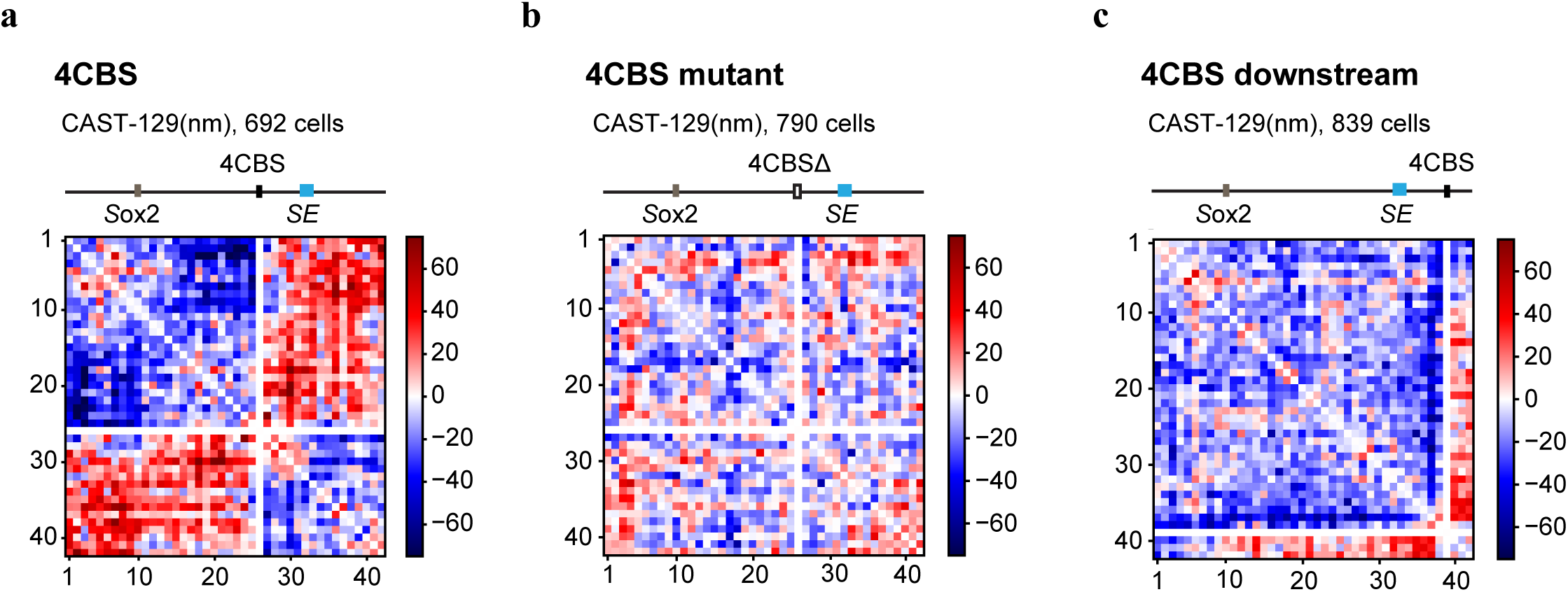
Radius of gyration of sub-domains. **a**, Difference of the median distance matrices between the CAST and 129 allele of the “4CBS” clone. **b**, Difference of the median distance matrices between the CAST and 129 allele of the “4CBS mutant” clone. **c**, Difference of the median distance matrices between the CAST and 129 allele of the “4CBS downstream” clone.

